# Breadboard Neuroscience: Exploring Neuronal Dynamics and Computation with Simple Electronic Circuits

**DOI:** 10.64898/2026.10.12.758658

**Authors:** Jack A. Supple

**Affiliations:** The Grass Fellowship Laboratory, Marine Biological Laboratory, Woods Hole, MA 02543, USA; Department of Bioengineering, Imperial College London, Exhibition Road, London, SW7 2AZ, UK

## Abstract

Simple electronic circuits provide a tangible means of exploring the electrical principles underlying neuronal signalling and computation. Here is presented a sequence of experiments using inexpensive, readily available components assembled on solderless breadboards to investigate passive membrane properties and analogue neural computation. A resistor-capacitor membrane model demonstrates how membrane resistance, capacitance, and excitatory conductance determine response amplitude and charging and discharging time constants. Multiple inputs illustrate sublinear summation and shunting inhibition. Connecting membrane compartments through axial resistances reproduces spatial attenuation and temporal delay. We then construct an elementary motion detector to demonstrate how simple electrical operations can produce direction-selective responses to visual motion. Together, this approach offers an accessible and flexible platform for exploring neuronal biophysics, allowing students to relate mathematical descriptions to observable electrical behaviour whilst developing practical electronics skills and engaging creatively in the design of their own experiments.

## Introduction

A fundamental feature of neuronal information processing is the generation, integration, and propagation of electrical signals, the dynamics of which emerge from the underlying biophysical properties of neurons. Membrane capacitance and ion-channel conductances, together with intracellular axial resistance, govern how electrical signals develop and propagate through neurons (Koch, 1999). These relationships are commonly expressed through equivalent electrical circuits and differential equations. For students of neuroscience unfamiliar with electronics, however, these formulations can often appear abstract and unintuitive. Constructing and manipulating physical circuits allows students to explore these relationships directly by varying individual component values and measuring the resulting voltage responses.

A range of educational tools have been developed to help students explore the electrical principles underlying neuronal function. Software such as MetaNeuron enables interactive exploration of membrane potentials, time constants, cable properties, and synaptic responses (Newman and Newman, 2013). Low-cost electrophysiological acquisition devices such as the SpikerBox make hands-on electrophysiology accessible to the classroom (Marzullo and Gage, 2012). Electronic neuron models offer another approach, using physical devices to reproduce key properties of neuronal activity. For example, Spikeling uses a microcontroller to simulate spiking-neuron dynamics in real time, while interfacing with external sensory stimuli and electronic synaptic inputs (Baden *et al*., 2018).

Physical analogue circuits provide a complementary approach, in which neuronal dynamics emerge directly from the electrical properties of circuit components. Dabrowski *et al*. (2013) described an undergraduate laboratory exercise in which students construct an electrical model of the cell membrane using a resistor and capacitor and examine how varying their values affects the resulting voltage response (Dabrowski *et al*., 2013). Giglia *et al*. (2019) presented a similar platform using 3D-printed component holders and Arduino-based measurements to explore passive membrane behaviour (Giglia *et al*., 2019). More recently, the Lu.i platform introduced an analogue implementation of a leaky integrate-and-fire neuron on a custom printed circuit board (PCB), enabling the exploration of spiking dynamics and interactions between neurons (Stradmann *et al*., 2025).

These approaches demonstrate the value of physical electronic circuits for exploring neuronal dynamics, but often rely on custom-fabricated components or printed circuit boards (PCBs), which may introduce additional fabrication requirements or limit the ease with which circuits can be modified and extended. In contrast, solderless breadboards offer an accessible alternative, allowing circuits to be constructed from inexpensive, readily available components without soldering or custom fabrication. This approach also introduces students to the fundamentals of electronic circuit design, an increasingly important skill in modern neuroscience and biological research more broadly.

Here, we present a sequence of breadboard neuroscience experiments that illustrate fundamental principles of neuronal dynamics and computation. Whilst these circuit dynamics arise directly from the electrical properties of the components, we incorporate a microcontroller to supply power and acquire voltage measurements, providing students with an additional introduction to microcontroller-based electronics. Part 1 develops a resistor–capacitor membrane model, introduces excitatory and shunting synaptic conductances, and extends the circuit into a multicompartment cable to examine spatial attenuation and temporal delay. Part 2 constructs an analogue implementation of the elementary motion detector (Borst and Helmstaedter, 2015). By incorporating light-dependent resistors, temporal filters, diode coincidence circuits, and an operational-amplifier subtractor, the circuit demonstrates how direction-selective responses to visual motion can emerge from elementary electrical operations.

## Part 1: Passive Membrane Dynamics

### 1.1 The Passive Membrane as an RC Circuit

The electrical properties of a cell membrane can be approximated as a resistor-capacitor (RC) circuit (Curtis and Cole, 1938; Fricke, 1925; Koch, 1999). This representation arises directly from the physical structure of the membrane. The phospholipid bilayer is electrically insulating and separates conductive intracellular and extracellular solutions. When a voltage difference exists across the membrane, ions redistribute according to their charge, but cannot readily cross the bilayer, causing opposite charges to accumulate on its two surfaces. This separation and storage of charge therefore causes the membrane to behave electrically as a capacitor. Ion channels provide finite conductive pathways through the otherwise insulating membrane and can therefore be represented electrically as resistances, or equivalently as conductances.

In the simplest passive-membrane model, the membrane capacitance, *C*_*m*_, is connected in parallel with an effective membrane resistance, *R*_*m*_ (Fig. 1A). The voltage across these components represents the membrane potential, *V*_*m*_. In a biological membrane, *R*_*m*_ represents the effective combined resistance of ion channels that are open around the resting potential, thereby providing a pathway for charge to leak across the otherwise insulating membrane. The associated reversal potential, *E*_*L*_, represents the voltage at which the net current through these resting ‘leak’ channels is zero.

**Figure 1:**
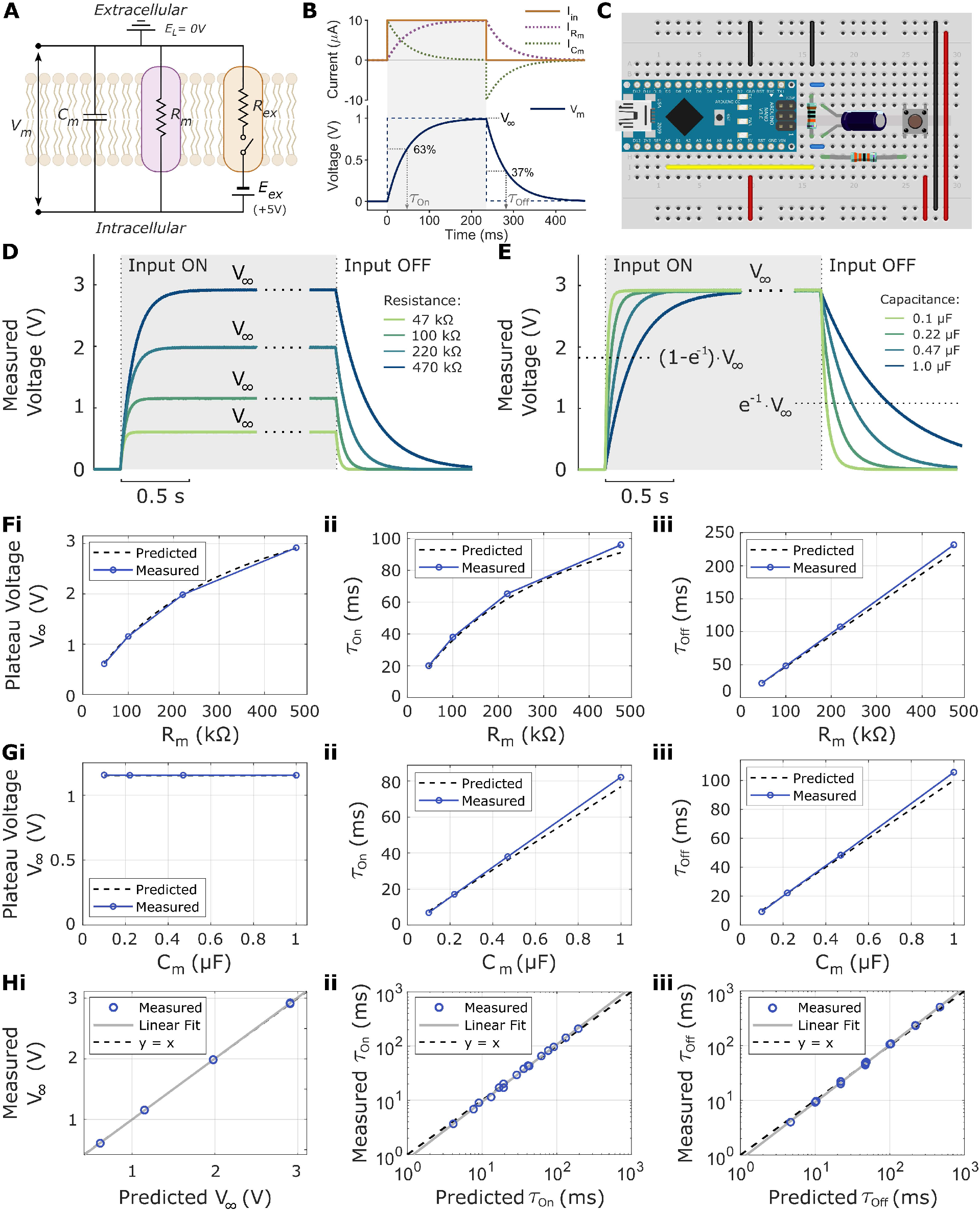
The passive membrane as an RC circuit. **(A)** Simplified electrical representation of a biological membrane. The phospholipid bilayer behaves as a membrane capacitor, *C*_*m*_, while resting ion conductances through open ion channels are represented by the membrane resistance, *R*_*m*_. Their associated ionic concentration gradients and membrane permeabilities determine the resting reversal potential, *E*_*L*_. The membrane voltage, *V*_*m*_, is measured across the membrane between the intracellular and extracellular compartments. An excitatory input is represented by an additional resistance, *R*_*ex*_, connected to its reversal potential, *E*_*ex*_; closing the switch activates this input. Note that in the present breadboard model, *E*_*L*_ = 0 V and *E*_*ex*_ = +5 *V*, such that membrane voltages are expressed relative to the resting potential. **(B)** Predicted voltage and current responses of a passive RC membrane to a square current input, *I*_*in*_. At input onset, current initially charges the membrane capacitor. As *V*_*m*_ approaches its steady-state value, *V*_∞_, capacitive current (*I*_*C*_) decreases while resistive current (*I*_*R*_) increases. After one membrane time constant, *τ*_*m*_, the voltage has reached 1 − *e*^−1^, or approximately 63%, of *V*_∞_. Following input offset, the capacitor discharges through the membrane resistance, with *V*_*m*_ falling to *e*^−1^, or approximately 37%, of its value at input offset after one time constant. Note that this illustrates the case of a constant-current input. In the present breadboard model, the input is instead supplied through a resistance, *R*_*ex*_, such that the input current decreases as *V*_*m*_ approaches the input reversal potential, *E*_*ex*_. The additional input conductance also reduces the effective membrane resistance during charging, resulting in a shorter charging time constant, *τ*_*on*_, than the discharging time constant, *τ*_*off*_. **(C)** Breadboard implementation of the passive membrane model. A capacitor (0.47 μF) and membrane resistor (100 kΩ) are connected in parallel, and a 5 V input is applied through a fixed 330 kΩ series resistor using a push button. Power is supplied from an Arduino Nano, and membrane voltage is sampled through its analogue input pin *A*_0_. Circuit ground represents the resting membrane potential, *E*_*L*_ = 0 V. The Arduino Nano serves only to supply power and measure the membrane voltage, without influencing the circuit dynamics. **(D)** Representative voltage responses obtained while varying *R*_*m*_ from 47 to 470 kΩ, with capacitance fixed at 0.47 *μ*F. Increasing *R*_*m*_ increases both the steady-state voltage, *V*_∞_, and the charging and discharging time constants. The long steady-state interval between the displayed onset- and offset-aligned windows was omitted and indicated with dotted lines. **(E)** Representative responses obtained while varying capacitance from 0.1 to 1.0 *μ*F, with *R*_*m*_ fixed at 470 kΩ. Increasing capacitance slows charging and discharging without changing the steady-state voltage. Dotted horizontal lines indicate *V*(*t* = *τ*_on_) = 0.63*V*_∞_, and *V*(*t* = *τ*_off_) = 0.37*V*_∞_, as indicated. Note that in both D and E, *τ*_on_ < *τ*_off_ due to the lower effective parallel resistance of *R*_*m*_and *R*_in_. **(F)** Measured (i) steady-state plateau voltage, *V*_∞_, (ii) charging time constant, *τ*_on_, and (iii) discharging time constant, *τ*_*off*_, as functions of membrane resistance, *R*_*m*_, using *C*_*m*_ = 0.47 μF, and *R*_*in*_ = 330 *kΩ*. Dashed lines indicate theoretical predictions. **(G)** Measured (i) steady-state plateau voltage, *V*_∞_, (ii) charging time-constant, *τ*_on_, and (iii) discharging time-constant, *τ*_*off*_, as functions of membrane capacitance, *C*_*m*_, using *R*_*m*_ = 100 kΩ, and *R*_*in*_ = 330 *kΩ*. Dashed lines indicate theoretical predictions. **(H)** Measured (i) steady-state plateau voltage, *V*_∞_, (ii) charging time constant, *τ*_*on*_, and (iii) discharging time constant, *τ*_*off*_, compared with their predicted values. The solid grey line shows the linear fit of the data (blue), and the dashed line shows the equality, *y* = *x*.

### 1.2 Charging and Discharging the Passive Membrane

The dynamics of the passive membrane arise from the time-dependent redistribution of current between the capacitive and resistive branches of the RC circuit (Fig. 1B). When current flows across the membrane, for example through an open ion channel, charge initially accumulates on *C*_*m*_, causing *V*_*m*_ to change according to

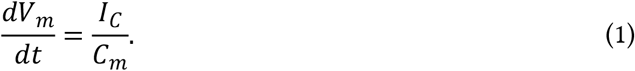

As *V*_*m*_ moves away from *E*_*L*_, the voltage across *R*_*m*_ increases in magnitude and a progressively larger fraction of the input current flows through the resistive pathway. Consequently, less current remains to charge *C*_*m*_, reducing the rate of change of *V*_*m*_. At the same time, *V*_*m*_ moves towards the input voltage, *V*_*in*_, reducing the voltage difference across *R*_in_. The input current is therefore

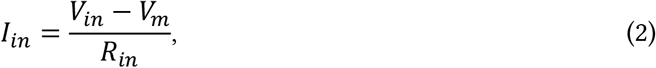

such that its magnitude decreases as *V*_*m*_ approaches *V*_*in*_. Kirchhoff’s current law requires that the current entering the membrane node equals the sum of the capacitive and resistive currents:

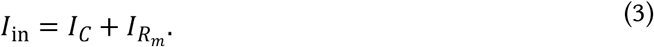

Substituting the current–voltage relationships for each element in equation 3 gives the first-order linear differential equation

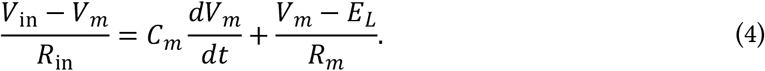

At steady state, *V*_*m*_ no longer changes, such that *dV*_*m*_/*dt* = 0. The steady-state membrane voltage is therefore

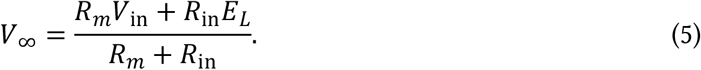

In the ground-referenced circuit used here, *E*_*L*_ = 0, giving

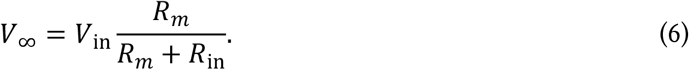

Thus, while the input is connected, *V*_*m*_ approaches *V*_∞_ *exponentially*, rather than approaching *V*_in_. Notably, the steady state *V*_∞_ is determined by both *R*_*m*_ and *R*_in_. Solving Eq. 4 gives the time course of this response:

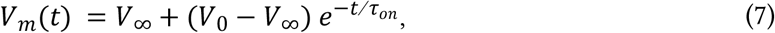

where *V*_0_ is the membrane voltage at input onset and

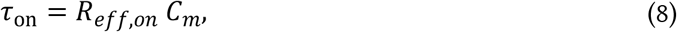

where *R*_*eff,on*_ is the effective parallel resistance of *R*_*m*_ and *R*_*in*_:

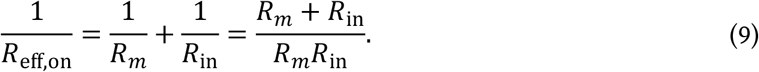

The time constant of charging, *τ*_on_, therefore depends on *C*_m_, *R*_m_, and *R*_in_ (a relationship we explore further in section 1.5). For the present case where *V*_0_ = *E*_*L*_ = 0, after one charging time constant,

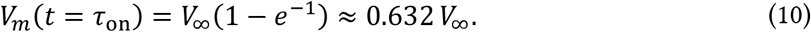

When the input branch is disconnected at *t* = *t*_end_, the capacitor discharges through *R*_*m*_, and *V*_*m*_ returns exponentially towards *E*_*L*_:

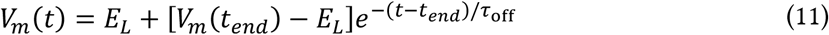

where

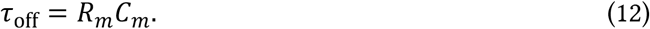

For *E*_*L*_ = 0, this reduces to

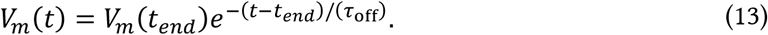

Therefore, in contrast to the charging time constant, the discharging time constant depends only on *R*_m_ and *C*_m_.

In summary, membrane resistance and capacitance influence passive membrane dynamics in distinct ways. For a given resistive voltage input, a high *R*_*m*_ reduces leak current, allowing a larger voltage to develop (Eq. 6), while also increasing the time required for the capacitor to charge or discharge (Eq. 8-9; Fig. 1D). Decreasing *R*_*m*_ has the opposite effects, with the voltage change becoming smaller and faster because current flows more readily through the leak pathway. Capacitance primarily affects response speed, as increasing *C*_*m*_ increases both *τ*_on_ and *τ*_off_, but does not alter *V*_∞_ (Fig. 1E).

### 1.3 Breadboard Implementation of the Passive Membrane

We can construct a breadboard model of the passive-membrane RC circuit using a resistor and capacitor connected in parallel (Fig. 1C; Fig. S1). An Arduino Nano provides a 5 V supply rail and provides a convenient way to measure *V*_*m*_ via analogue input A0 (Fig. 1C, yellow). In this breadboard passive membrane model, circuit ground is defined as the resting potential, such that *E*_*L*_ = 0 V. Voltages in the model therefore represent changes relative to rest rather than literal physiological membrane voltages. Furthermore, whilst the absolute voltages are orders of magnitude different from those across biological membranes, we can nonetheless observe the same dynamic phenomena as those seen in real neurons.

A push button provides a resistive voltage input (*V*_in_) by connecting *V*_*m*_ to the 5 V rail through an input resistor (here 330 kΩ; Fig. 1C). The resulting input current, *I*_in_, depends on the instantaneous driving force across the resistor according to Eq. 2, and is therefore analogous to a conductance-based synaptic input. The input resistor (*R*_in_) limits the current available to charge *C*_*m*_, allowing *V*_*m*_ to rise gradually towards its steady-state value.

### 1.4 Observing the Effects of Membrane Resistance and Capacitance on Passive Dynamics

The separate effects of *R*_*m*_ and *C*_*m*_ can readily be observed on the breadboard model (Fig. 1D-G). Varying resistance changes both response amplitude and kinetics (Fig. 1D,F), whereas varying capacitance changes the time-course of charging and discharging, while leaving *V*_∞_ unchanged (Fig. 1E,G). The measured steady state voltage, *V*_∞_, closely matched the expected value across a range of tested resistor and capacitor combinations (Fig. 1Hi; *R*^2^ = 0.999). Similarly, the measured charging and discharging time constants closely followed the theoretical RC predictions (*τ*_on_ followed *R*_*eff,on*_ *C*_*m*_ with *R*^2^ = 0.998, Fig. 1Hii; *τ*_off_ followed *R*_*m*_ *C*_*m*_ with *R*^2^ = 0.996, Fig. 1Hiii). These results establish that a small number of inexpensive breadboard components can demonstrate the central passive electrical properties of a biological membrane, including resistance-dependent determination of steady-state membrane voltage, charge storage by the capacitive lipid bilayer, and resistance- and capacitance-dependent membrane time constants.

### 1.5 Synaptic Inputs as Conductances

Let us further consider the biological analogy of applying external input to the membrane RC circuit. In sections 1.2–1.4, current (*I*_in_) was introduced to the circuit by connecting *V*_*m*_ to the 5 V rail (*V*_in_) through a push button and input resistor, *R*_in_. Electrically, this is equivalent to opening an ohmic synaptic conductance connecting the membrane to a synaptic reversal potential. In the present circuit, we reinterpret this as an excitatory synaptic input, such that

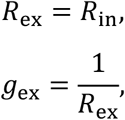

and the 5 V supply rail represents the excitatory reversal potential *E*_ex_. The resulting synaptic current depends on the instantaneous driving force between the synaptic reversal potential and the membrane potential:

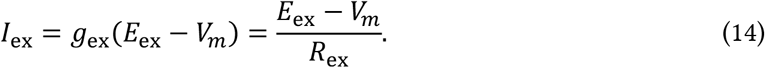

The button therefore produces a step increase in conductance rather than a constant-current input. As *V*_*m*_ approaches *E*_ex_, the driving force decreases and the synaptic current becomes smaller. The effects of synaptic conductance can be examined on the breadboard by holding *R*_*m*_ and *C*_*m*_ constant while varying *R*_ex_ (Fig. 2). From Eq. 6, using *E*_*L*_ = 0, and *R*_in_ = *R*_*ex*_, the steady-state membrane voltage is

**Figure 2:**
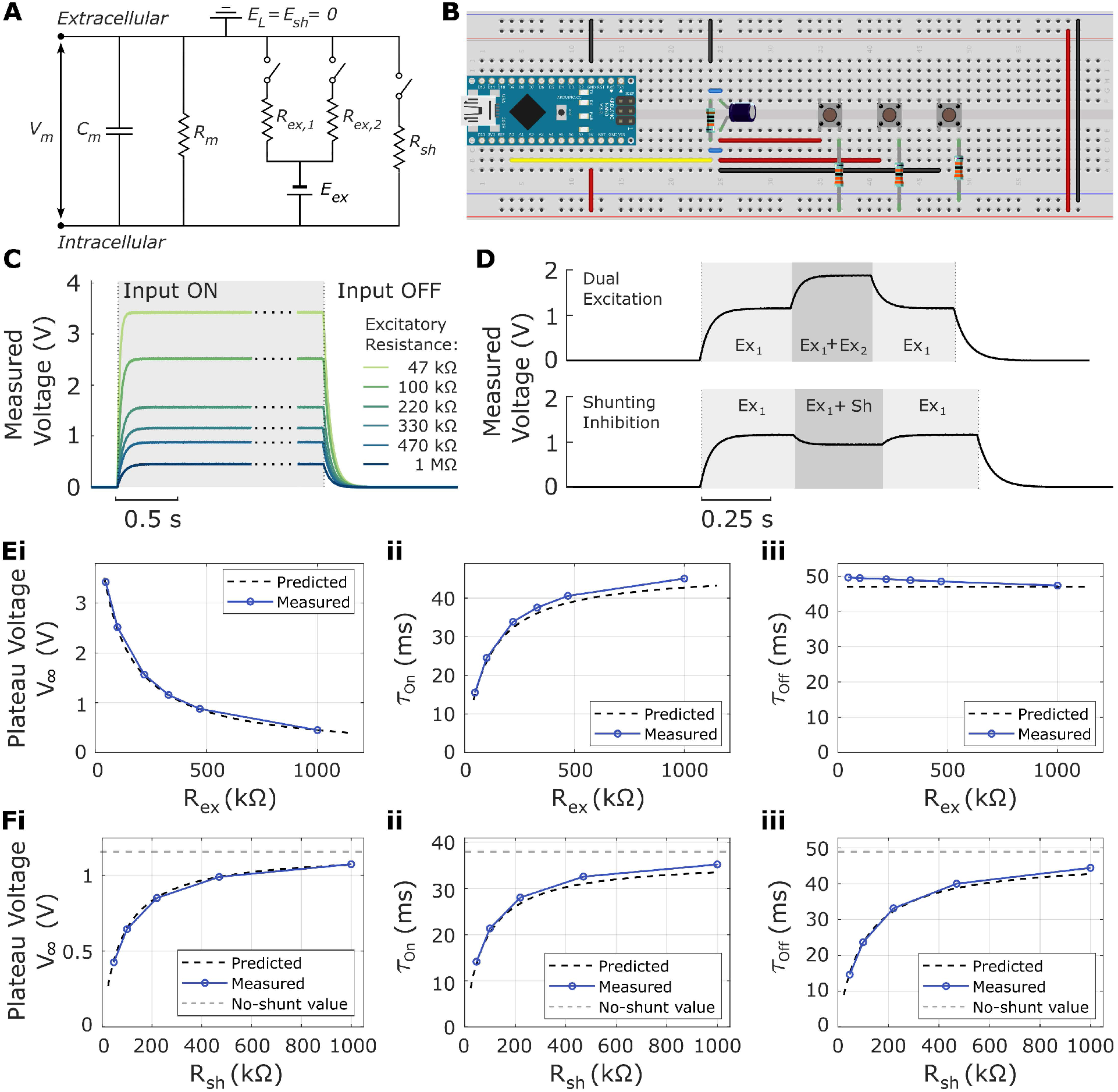
Excitatory and Inhibitory Synaptic Conductances. **(A)** Circuit diagram of the passive membrane, comprising membrane capacitance, *C*_*m*_, and membrane resistance, *R*_*m*_, connected to the leak reversal potential, *E*_*L*_ = 0. Excitatory inputs are delivered from *E*_*ex*_ through switchable excitatory resistances, *R*_*ex*_, while a switchable shunt resistance, *R*_*sh*_, provides an additional conductance to *E*_*L*_. (**B**) Breadboard implementation of the circuit, with membrane voltage measured using an Arduino Nano. Here *E*_*L*_ = 0 *V*, and *E*_*ex*_ = +5 *V*. Two buttons connect the membrane node to 5 V via resistors *R*_*ex*_ = 330 *kΩ* each. A third button provides a shunting pathway to circuit ground via resistor *R*_*sh*_ = 330 *kΩ*. **(C)** Representative membrane-voltage responses to excitation through different values of *R*_*ex*_. Decreasing *R*_*ex*_ increases excitatory conductance, producing a larger steady-state depolarisation and a faster voltage rise following input onset. Following input offset, the membrane returns towards *E*_*L*_ through *R*_*m*_. **(D)** Representative membrane-voltage responses to multiple simultaneous inputs. Top, activation of a second excitatory input (*Ex*_1_ + *Ex*_2_) produces additional depolarisation relative to *Ex*_1_ alone. Bottom, activation of the shunting conductance (*Ex*_1_ + *Sh*) reduces the depolarisation produced by *Ex*_1_. Shaded regions indicate the periods during which the additional input is active. In this example, the shunting conductance is activated transiently during excitation and removed before the excitatory input ends. **(E)** Effects of excitatory resistance, *R*_*ex*_, on **(i)** steady-state plateau voltage, *V*_∞_, **(ii)** membrane charging time constant, *τ*_*on*_, and **(iii)** membrane discharging time constant, *τ*_*off*_. Increasing *R*_*ex*_ reduces excitatory conductance and therefore decreases *V*_∞_ while increasing *τ*_*on*_. In contrast, *τ*_*off*_ remains independent of *R*_*ex*_, because the excitatory conductance is removed following input offset. Dashed black lines show theoretical predictions and blue points show measured values. **(F)** Effects of shunt resistance, *R*_*sh*_, on **(i)** *V*_∞_, **(ii)** *τ*_*on*_, and **(iii)** *τ*_*off*_ during shunting inhibition. For these measurements, the shunting conductance remains connected throughout excitation and the subsequent discharge, allowing its effects on both *τ*_on_ and *τ*_off_ to be measured. Decreasing *R*_*sh*_ increases the shunting conductance, reducing *V*_∞_ and shortening both *τ*_*on*_ and *τ*_*off*_. Dashed black lines show theoretical predictions, blue points show measured values, and grey dashed lines indicate the corresponding values in the absence of the shunt.

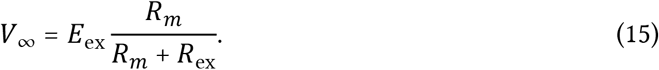

Increasing *R*_ex_ reduces the synaptic conductance and therefore reduces the current entering the membrane for a given driving force. A larger fraction of the voltage difference between *E*_ex_ and *E*_*L*_ is consequently dropped across *R*_ex_, and *V*_∞_ remains closer to the resting potential. Conversely, decreasing *R*_ex_ produces a stronger synaptic conductance and moves *V*_∞_ closer to *E*_ex_. This trend can be seen on the breadboard model in Fig. 2Ei, with *V*_∞_ approaching 0 as excitatory resistance increases.

The synaptic resistance also affects the charging time constant according to Eqs. 8-9:

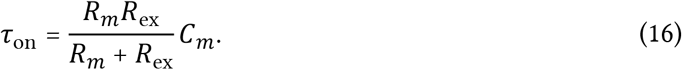

As *R*_*ex*_ increases, the synaptic conductance decreases, reducing the additional conductive pathway across the membrane. The combined resistance through *R*_*m*_ and *R*_ex_ therefore increases, increasing the charging time constant *τ*_on_. Conversely, a low *R*_ex_ increases the combined membrane conductance and shortens *τ*_on_. This relationship between *R*_ex_ and *τ*_on_ can be seen on the breadboard model (Fig. 2Eii).

Conductance-based excitation therefore affects both the amplitude and time course of membrane charging. In contrast, the passive discharging time constant, *τ*_off_, is independent of *R*_*ex*_ because the synaptic branch is disconnected during discharge. This independence of *τ*_off_ from *R*_*ex*_ is closely replicated on the breadboard (Fig. 2Eiii).

### 1.6 Multiple Synaptic Inputs

We can next examine the integration of multiple synaptic inputs. Identical inputs connected to the same reversal potential act as parallel conductances. For *n* inputs, each with resistance *R*_ex_, the effective excitatory resistance, *R*_ex,eff_, decreases as

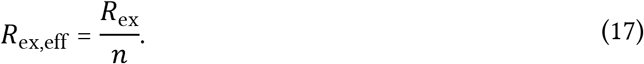

Substituting *R*_ex,eff_ for *R*_in_, and *E*_ex_ for *V*_in_, in Eq. 5 gives the steady-state membrane voltage

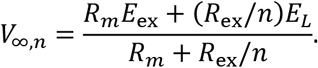

For the ground-referenced circuit, where (*E*_L_=0), this reduces to

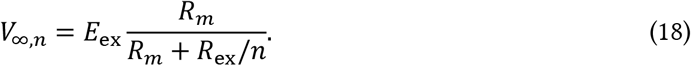

Therefore, recruiting additional identical synaptic conductances progressively reduces the effective input resistance, *R*_ex,eff_, allowing *V*_*m*_ to move closer to *E*_ex_. Importantly, as *V*_*m*_ approaches *E*_*ex*_, the driving force, *E*_ex_ − *V*_*m*_, decreases. Each additional identical input therefore produces a progressively smaller increment in steady-state membrane voltage, resulting in sublinear summation, as observed in the breadboard model (Fig. 2D).

Because adding identical excitatory inputs is mathematically equivalent to reducing the overall excitatory resistance, its effects on *V*_∞_, and *τ*_on_ follow directly from the relationships established for a single excitatory input in Fig. 2E. Recruiting additional inputs increases *V*_∞_ while simultaneously increasing the total membrane conductance and shortening *τ*_on_. Once the excitatory inputs are removed, however, the passive *τ*_off_ remains determined by *R*_*m*_*C*_*m*_, and is therefore independent of the number of previously active inputs, just as it is independent of *R*_*ex*_ (Fig. 2Eiii).

Thus, the breadboard model of multiple synaptic inputs (Fig. 2B; Fig. S2) demonstrates how the recruitment of additional synaptic conductances simultaneously alters both the amplitude and kinetics of the membrane response.

### 1.7 Depolarisation, Hyperpolarisation, and Shunting Inhibition

The same electrical principles also apply to inhibitory synaptic inputs. As described above, the excitatory conductance, *g*_ex_ = 1/*R*_ex_, connects the membrane to an excitatory reversal potential, *E*_ex_, lying above the resting membrane potential, *E*_*L*_. An inhibitory input can be represented analogously by a conductance, *g*_inh_ = 1/*R*_inh_, connecting the membrane to an inhibitory reversal potential, *E*_inh_. Whether this input produces hyperpolarising or shunting inhibition depends on the position of *E*_inh_relative to the resting membrane potential.

Hyperpolarising inhibition occurs when *E*_inh_ < *E*_*L*_, whereas shunting inhibition occurs when *E*_inh_ ≈ *E*_*L*_. Because circuit ground (0 V) represents *E*_*L*_ in the present breadboard model, voltages below *E*_*L*_ cannot be represented without first offsetting the resting potential above ground. The same circuit can, however, demonstrate shunting inhibition.

Consider a shunting conductance *g*_*sh*_ = 1/*R*_*sh*_ with

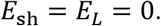

The current through this conductance is

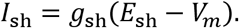

At rest, *V*_*m*_ = *E*_*L*_ = *E*_sh_, and therefore *I*_sh_ = 0. Opening the shunting conductance consequently produces little or no change in membrane voltage by itself, distinguishing shunting inhibition from hyperpolarising inhibition. However, the effect becomes apparent when another input depolarises *V*_*m*_ (Fig. 2D). Once *V*_*m*_ moves above *E*_sh_, a driving force develops across the shunting conductance. The shunt therefore opposes the depolarisation and provides an additional conductive pathway back towards *E*_*L*_.

In the breadboard circuit (Fig. 2B; Fig. S2), *R*_*m*_ and *R*_*sh*_ both connect *V*_*m*_ to ground in parallel, so the addition of *R*_*sh*_ can be thought of as lowering the effective membrane resistance. Because *R*_*m*_ and *R*_sh_ share the same reversal potential in this circuit, adding *R*_sh_ is electrically equivalent to reducing the effective resistance between *V*_*m*_ and *E*_*L*_, and therefore produces the same qualitative effects as reducing *R*_*m*_ (Fig. 1F). The addition of *R*_sh_ reduces *V*_∞_, with smaller *R*_sh_ creating a larger reduction in *V*_∞_ (Fig. 2Fi). Similarly, the addition of *R*_sh_ shortens the charging time constant, *τ*_on_, with greater reductions occurring with smaller *R*_sh_ values (Fig. 2Fii).

Notably, *R*_sh_ also affects the discharging time constant, *τ*_off_, because the shunting pathway remains connected after the excitatory input is removed. During discharge, *C*_*m*_ therefore discharges through *R*_*m*_ and *R*_sh_ in parallel, giving

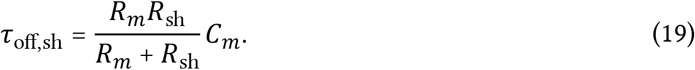

Consequently, smaller *R*_*sh*_ values produce faster membrane discharge (Fig. 2Fiii), in contrast to the independence of τ_off_ from *R*_*ex*_ when the excitatory branch is disconnected (Fig. 2Eiii).

### 1.8 Passive Voltage Spread Along a Cable

The passive membrane model described above treats the membrane as a single electrical compartment. However, dendrites and axons are spatially extended, so a voltage change generated at one location must spread through the intracellular medium to influence more distant regions. A simple passive cable can be represented by connecting multiple RC membrane compartments through an axial (cytoplasmic) resistance, *R*_*a*_ (Fig. 3A). Each compartment contains the same membrane resistance, *R*_*m*_, and capacitance, *C*_*m*_, while *R*_*a*_ represents the resistance to current flow along the interior of the neurite. This arrangement can be reproduced directly on a breadboard by constructing a chain of identical RC compartments linked by axial resistors, with the voltage at each compartment measured independently (Fig. 3B; Fig. S3).

**Figure 3:**
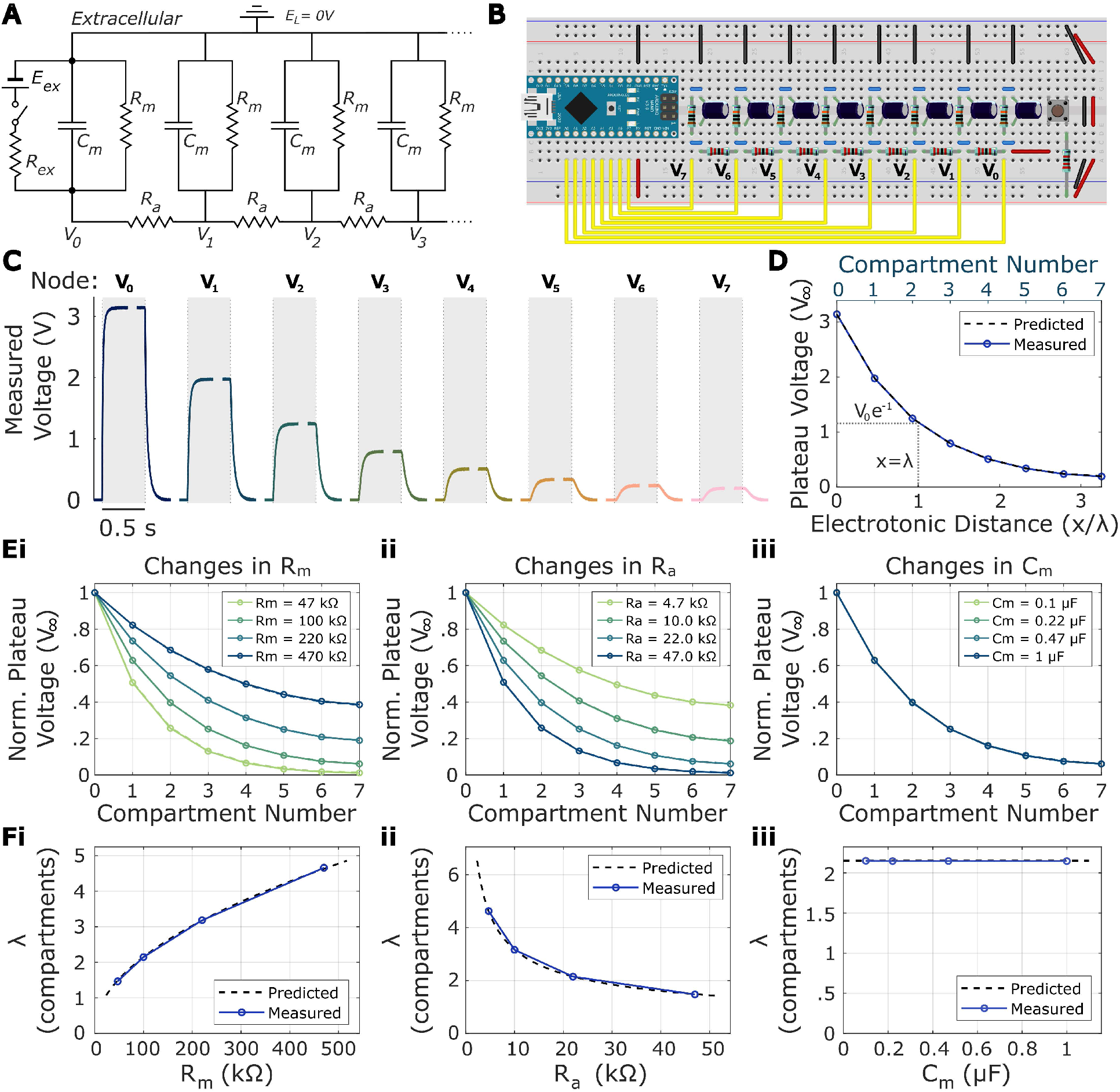
Spatial properties of the passive cable model. **(A)** Schematic of the passive cable model, comprising membrane compartments with membrane resistance *R*_*m*_ and capacitance *C*_*m*_ connected in parallel to the leak reversal potential, *E*_*L*_ = 0, and coupled to neighbouring compartments by axial resistances, *R*_*a*_. Excitatory input is delivered to node *V*_0_ through the excitatory resistance *R*_*ex*_, while neighbouring compartments are coupled by the axial resistance *R*_*a*_. Membrane voltages are measured at the corresponding compartment nodes, *V*_0_ − *V*_7_. **(B)** Breadboard implementation of the eight-compartment cable, with membrane voltages *V*_0_ − *V*_7_ sampled by the Arduino Nano analog input pins A0-A7. The baseline cable parameters used here are *R*_*ex*_ = 22 *kΩ, R*_*m*_ = 100 *kΩ, R*_*a*_ = 22 *kΩ, C*_*m*_ = 0.47 *μF*. A lower *R*_*ex*_ was used compared to Figs. 1-2 to produce a larger depolarisation at the input node to facilitate visualisation of the voltage spread through the compartments. **(C)** Representative voltage responses recorded from *V*_0_ − *V*_7_ following excitation at *V*_0_. Grey shading indicates the period of excitation. Voltage amplitude progressively attenuates with distance from the input. **(D)** Steady-state plateau voltage *V*_∞_ as a function of electrotonic distance, *x*/*λ*, for the baseline cable (*R*_*m*_ = 100 *kΩ, R*_*a*_ = 22 *kΩ, R*_*ex*_ = 22 *kΩ, C*_*m*_ = 0.47 *μF*). The corresponding compartment numbers are shown on the upper axis. The dashed line shows the theoretical voltage profile calculated from the finite eight-compartment circuit. Electrotonic distance is calculated using the theoretical space constant of an infinite discrete cable. Dotted guides indicate a distance of one space constant, *λ*, at which the ideal exponential profile decays to 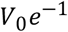. **(E)** Normalized steady-state plateau voltages, *V*_∞_, following variation of **(i)** *R*_*m*_, **(ii)** *R*_*a*_, and **(iii)** *C*_*m*_, with all other component values held at the baseline values used in C-D. Increasing *R*_*m*_ increases the spatial spread of voltage, whereas increasing *R*_*a*_ decreases it; changing *C*_*m*_ does not alter the steady-state spatial profile. **(F)** Space constant, *λ*, measured from the voltage profiles as a function of **(i)** *R*_*m*_, **(ii)** *R*_*a*_, and **(iii)** *C*_*m*_. Dashed lines show predictions from the passive cable model and blue points/lines show measured values. As predicted, *λ* increases with *R*_*m*_, decreases with *R*_*a*_, and is independent of *C*_*m*_.

When one compartment is depolarised, current can either spread longitudinally through *R*_*a*_ into neighbouring compartments, or leave through the local membrane resistance, *R*_*m*_. Consequently, some current is lost across the membrane at each successive compartment, causing the voltage response to decrease progressively with distance from the input (Fig. 3C-D).

For a given compartment i, we saw in Eq. 4 that incoming current equals the sum of capacitive and resistive currents. For an interior cable compartment, *i*, the membrane voltage *V*_*m*_ in Eq. 4 is replaced by the local compartment voltage *V*_*i*_. The single input-current term is also replaced by two axial current terms, representing current exchanged through *R*_*a*_ with the neighbouring compartments *i* − 1 and *i* + 1. Eq. 4 therefore becomes

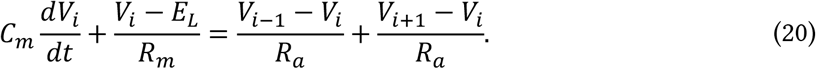

At steady state, membrane voltage is no longer changing, such that *dV*_*i*_/*dt* = 0 and the capacitive current is zero. In the breadboard model, the leak reversal potential is represented by electrical ground, so *E*_*L*_ = 0. Eq. 20 therefore reduces to:

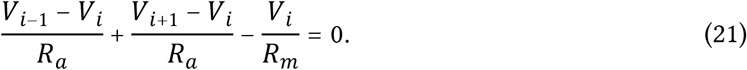

Rearranging:

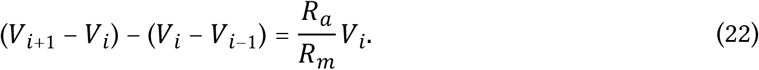

The two terms in parentheses on the left describe the voltage differences between compartment *i* and each of its two neighbouring compartments. The difference between these terms therefore describes how the voltage difference itself changes from one part of the cable to the next, and is known as the second difference. To express this change with respect to physical distance rather than compartment number, let neighbouring compartments be separated by a distance Δ*x*. Dividing each voltage difference by Δ*x* gives the local voltage gradient, and dividing the change in this gradient by Δ*x* gives

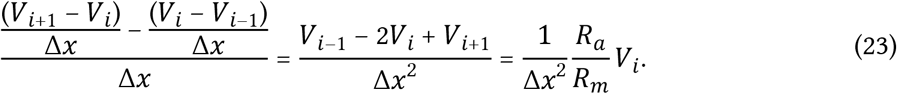

Note that *R*_*a*_ and *R*_*m*_ here refer to the compartmental axial and membrane resistances, respectively, and therefore depend on the length of each compartment, Δ*x*. Axial resistance increases with compartment length because current must travel through a longer stretch of cytoplasm, such that

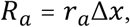

where *r*_*a*_ is the axial resistance per unit length. In contrast, membrane resistance decreases with compartment length because a longer compartment contains more membrane area and therefore more parallel pathways for current to leave the cable. Thus,

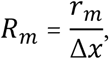

where *r*_*m*_ is the membrane resistance of a unit length of cable. Substituting these expressions into Eq. 23 gives

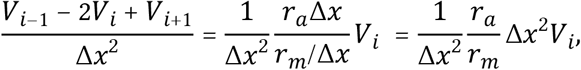

which simplifies to

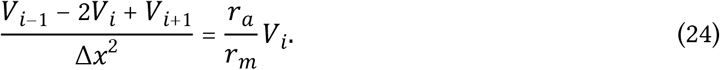

As Δx becomes progressively smaller, the discrete second difference term on the left approaches the continuous second spatial derivative, giving the steady-state cable equation:

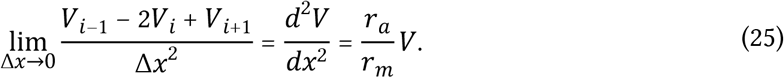

Eq. 25 shows that the spatial change in membrane voltage depends on the relative axial and membrane resistances of the cable. Increasing membrane resistance, *r*_*m*_, reduces current loss across the membrane and allows voltage to spread farther, whereas increasing axial resistance, *r*_*a*_, restricts longitudinal current flow and causes voltage to attenuate more rapidly. This relationship can be expressed as a characteristic length, known as the space constant, *λ*:

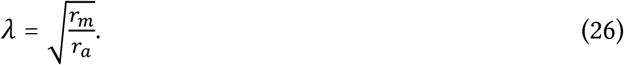

The steady-state cable equation therefore becomes

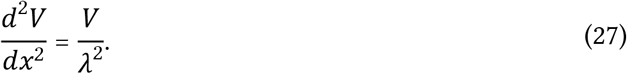

For a passive cable extending away from an input, voltage cannot increase without bound with distance, leaving the decaying solution

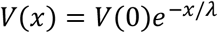

where *V*(0) is the voltage at the input location. At a distance of one space constant, *λ*:

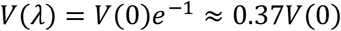

Thus, λ provides a convenient measure of the distance over which a passive voltage change spreads along the cable.

This spatial attenuation can be observed directly in the discretised breadboard cable model, where excitation of the first compartment produces progressively smaller voltage responses at successive compartments (Fig. 3C). At steady state, voltage decreased with electrotonic distance from the input, reaching approximately *V*_0_*e*^−1^, or 37% of the input voltage, at one space constant (Fig. 3D). Varying the values of *R*_*m*_, *R*_*a*_, and *C*_*m*_ demonstrates how the passive electrical properties of the cable determine the spatial spread of voltage. Increasing *R*_*m*_ reduces current loss across the membrane and therefore allows voltage to spread farther, whereas increasing *R*_*a*_ restricts axial current flow and reduces the distance over which voltage spreads (Fig. 3Ei–ii). In contrast, changing *C*_*m*_ does not alter the steady-state voltage profile because capacitive current is zero at steady state (Fig. 3Eiii). Accordingly, the measured space constant increases with *R*_*m*_, decreases with *R*_*a*_, and remains independent of *C*_*m*_, in agreement with the predictions of the passive cable model (Fig. 3F).

### 1.9 Propagation Delay Along a Cable

Passive voltage spread along a cable is not only attenuated with distance but also delayed (Fig. 4). Although each compartment has the same intrinsic membrane time constant, *τ*_*m*_ = *R*_*m*_*C*_*m*_, current must progressively charge the membrane capacitance of successive regions as it spreads along the cable. Consequently, more distant regions respond later than regions close to the input (Fig. 4Ai).

**Figure 4:**
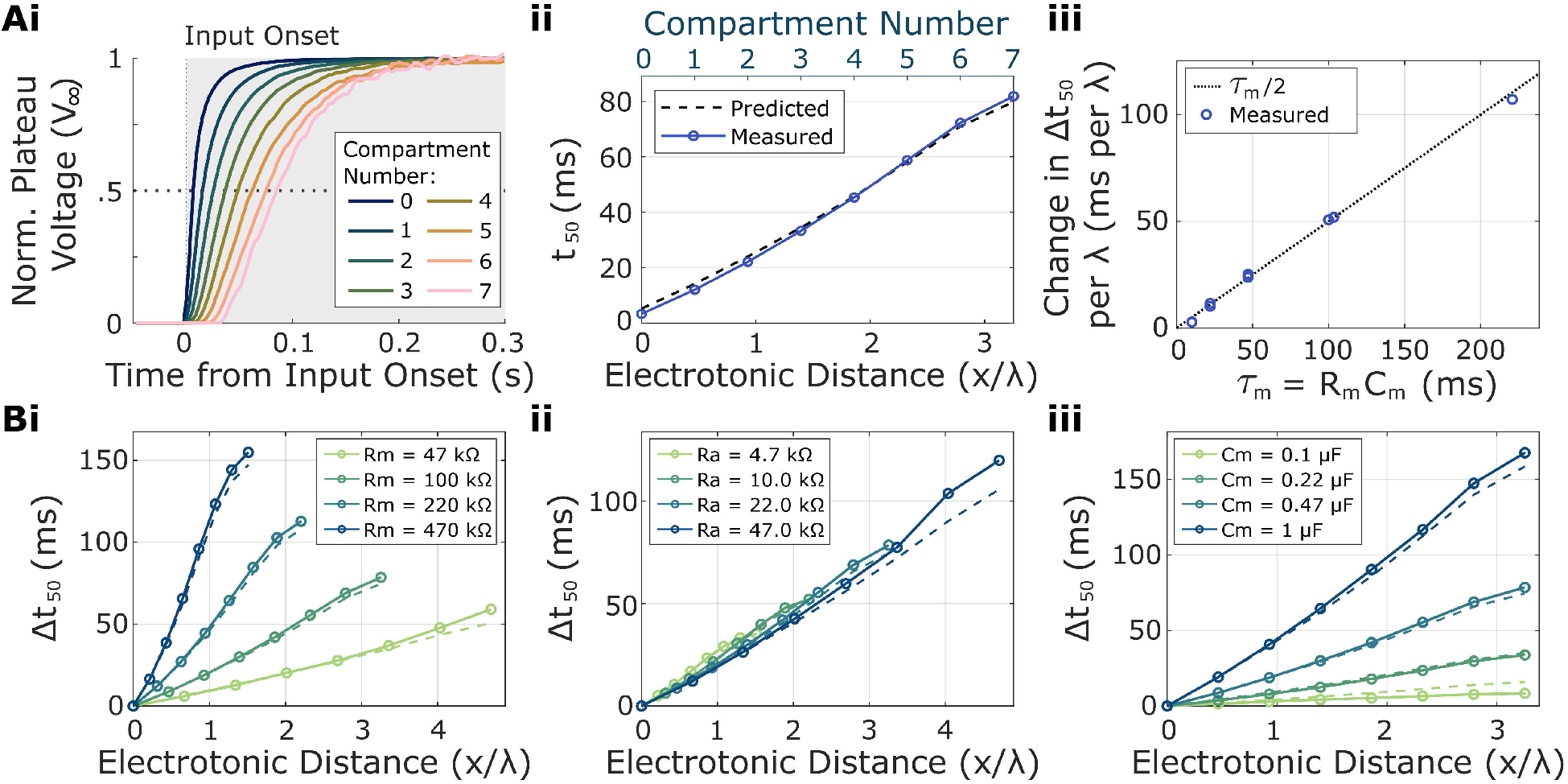
Temporal properties of passive voltage spread along the cable. **(A)** Temporal response of the baseline eight-compartment cable from Fig. 3 (*R*_*m*_ = 100 *kΩ, R*_*a*_ = 22 *kΩ, C*_*m*_ = 0.47 *μF*). **(i)** Normalized voltage responses of compartments *V*_0_ − *V*_7_, aligned to the onset of excitation. More distant compartments respond progressively later. **(ii)** Time required for each compartment to reach 50% of its local steady-state response, *t*_50_, plotted against electrotonic distance, *x*/*λ*. The upper axis indicates the corresponding compartment number. The dashed line shows the prediction of the finite eight-compartment cable model. **(iii)** Gradient of the relative *t*_50_ delay, Δ*t*_50_, per space constant, *λ*, plotted as a function of membrane time constant, *τ*_*m*_. The dotted line indicates the theoretical prediction of *τ*_*m*_/2 per space constant for centroid propagation delays in an infinite continuous cable, shown for comparison. **(B)** Relative propagation delay, Δ*t*_50_, as a function of electrotonic distance following systematic variation of (i) *R*_*m*_, (ii) *R*_*a*_, and (iii) *C*_*m*_, with all other component values held at their baseline values. Increasing *R*_*m*_ or *C*_*m*_ increases *τ*_*m*_ and produces a steeper increase in Δ*t*_50_ with electrotonic distance. Changing *R*_*a*_ alters the space constant, *λ*, but not *τ*_*m*_, causing the relationships to align when distance is expressed in electrotonic units. Solid points show breadboard measurements; dashed lines show predictions from the finite eight-compartment cable model.

The steady-state cable equation derived above describes the spatial distribution of voltage once membrane charging is complete. To describe how voltage changes over both space and time, the capacitive current must instead be retained, giving the time-dependent cable equation

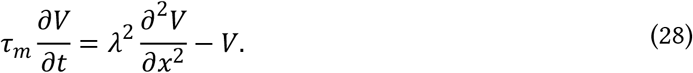

For a semi-infinite continuous cable, the centroid delay relative to the input increases by *τ*_*m*_/2 per space constant, giving

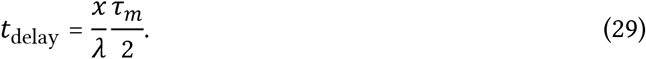

The distance, x, along the cable can also be expressed in units of the space constant by setting *x* = *nλ*, giving

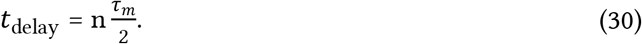

Thus, each additional space constant introduces a characteristic delay of *τ*_*m*_/2. Importantly, this does not mean that the local membrane time constant increases with distance. Each region of a uniform cable retains the same intrinsic *τ*_*m*_; the increasing delay instead arises from the progressive spread of current through, and charging of, the intervening membrane.

Whilst the characteristic delay is defined from the centre-of-mass of the voltage response derivative (Agmon-Snir and Segev, 1993; Koch, 1999), here we use the time required for each compartment to reach 50% of its local steady-state response, *t*_50_, as an intuitive measure of response timing (Fig. 4Aii). Subtracting the corresponding time at the input compartment gives the relative propagation delay:

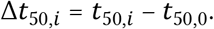

In the breadboard cable, Δ*t*_50_ increases approximately linearly with electrotonic distance (Fig. 4Aii), and the resulting delay gradient scales approximately with *τ*_*m*_/2 (Fig. 4Aiii).

Varying *R*_*m*_, *R*_*a*_, and *C*_*m*_ further demonstrates how the electrical properties of the cable determine this propagation delay (Fig. 4B). Increasing *R*_*m*_ increases the membrane time constant, *τ*_*m*_ = *R*_*m*_*C*_*m*_, and therefore increases the *t*_50_ delay accumulated per unit electrotonic distance according to Eq. 29 (Fig. 4Bi). In contrast, increasing *R*_*a*_ changes the space constant, and therefore the conversion between compartment number and electrotonic distance, but does not alter *τ*_*m*_. Consequently, the *R*_*a*_ conditions show similar relationships between Δ*t*_50_ and *x*/*λ* once distance is expressed electrotonically (Fig. 4Bii). Increasing *C*_*m*_ leaves the space constant unchanged but increases *τ*_*m*_, producing progressively steeper relationships between Δ*t*_50_ and electrotonic distance (Fig. 4Biii). Together, these results show that passive propagation delay is determined by both the electrotonic distance travelled and the intrinsic membrane time constant.

The breadboard cable model therefore demonstrates two consequences of passive cable propagation. With increasing distance, the amplitude of the steady-state voltage decreases according to the space constant, λ, while the temporal response becomes progressively delayed according to both electrotonic distance and the membrane time constant, *τ*_*m*_. Thus, passive cable properties determine not only how far a voltage signal spreads, but also how quickly it reaches more distant regions.

## Part 2: Computing in Analogue

The circuits introduced in Part 1 demonstrate how the passive electrical properties of neurons shape voltage in time and space. These passive electrical properties provide a physical substrate for neural computation, where signals with different amplitudes, spatial origins, and temporal dynamics can interact as they propagate through a circuit. Importantly, computation can already occur at the level of graded membrane potentials, without requiring the generation of action potentials. Inputs can be weighted by their conductance and integrated, filtered over time, and combined nonlinearly to transform information before any spike is produced.

To demonstrate how relatively simple electrical components can perform a biologically relevant computation, we will next construct an analogue model of an **elementary motion detector (EMD)**. Elementary motion detection compares signals from neighbouring photoreceptors across both space and time, allowing the circuit to distinguish the direction of image motion. This provides a classic example of neural computation because directional motion cannot be inferred from the activity of a single photoreceptor alone. Instead, neighbouring signals must be temporally filtered, combined nonlinearly, and compared through opponent subtraction to generate a direction-selective response.

### 2.1 The Elementary Motion Detector

Consider two neighbouring photoreceptors, A and B, viewing adjacent regions of visual space. A stationary change in illumination can activate either receptor, but the response of an individual receptor contains no information about the direction in which an image is moving. Direction becomes available only by comparing the timing of responses across space. If an image feature moves from A towards B, receptor A responds first, and receptor B responds shortly afterwards. Motion in the opposite direction produces the reverse sequence.

For a neuron to detect motion, it must distinguish these two temporal sequences. This requires the circuit to retain information about the earlier photoreceptor response long enough to compare it with the later response of its neighbour. A simple way to achieve this is to delay one signal and combine it with the instantaneous signal from the neighbouring receptor. If the delay approximately matches the time required for the image to move between the two receptors, the two signals overlap strongly for one direction of motion but less strongly for the opposite direction. This basic principle forms the foundation of the EMD.

Here, we focus on the classical multiplicative EMD formulated by Hassenstein and Reichardt (Hassenstein and Reichardt, 1956), which compares signals from two neighbouring points in visual space separated by a distance Δ*x* (Borst and Helmstaedter, 2015; Buchner, 1984). In one half-detector, the signal from photoreceptor A is temporally filtered before being multiplied by the instantaneous signal from photoreceptor B (Fig. 5Ai). Temporal filtering prolongs the A response, allowing it to overlap with the subsequent B response during motion from A to B. Multiplication produces a large output when both signals are simultaneously elevated, but a smaller output when either signal is low. The half-detector therefore responds more strongly to motion from A to B than to the reverse sequence, establishing its preferred direction.

**Figure 5:**
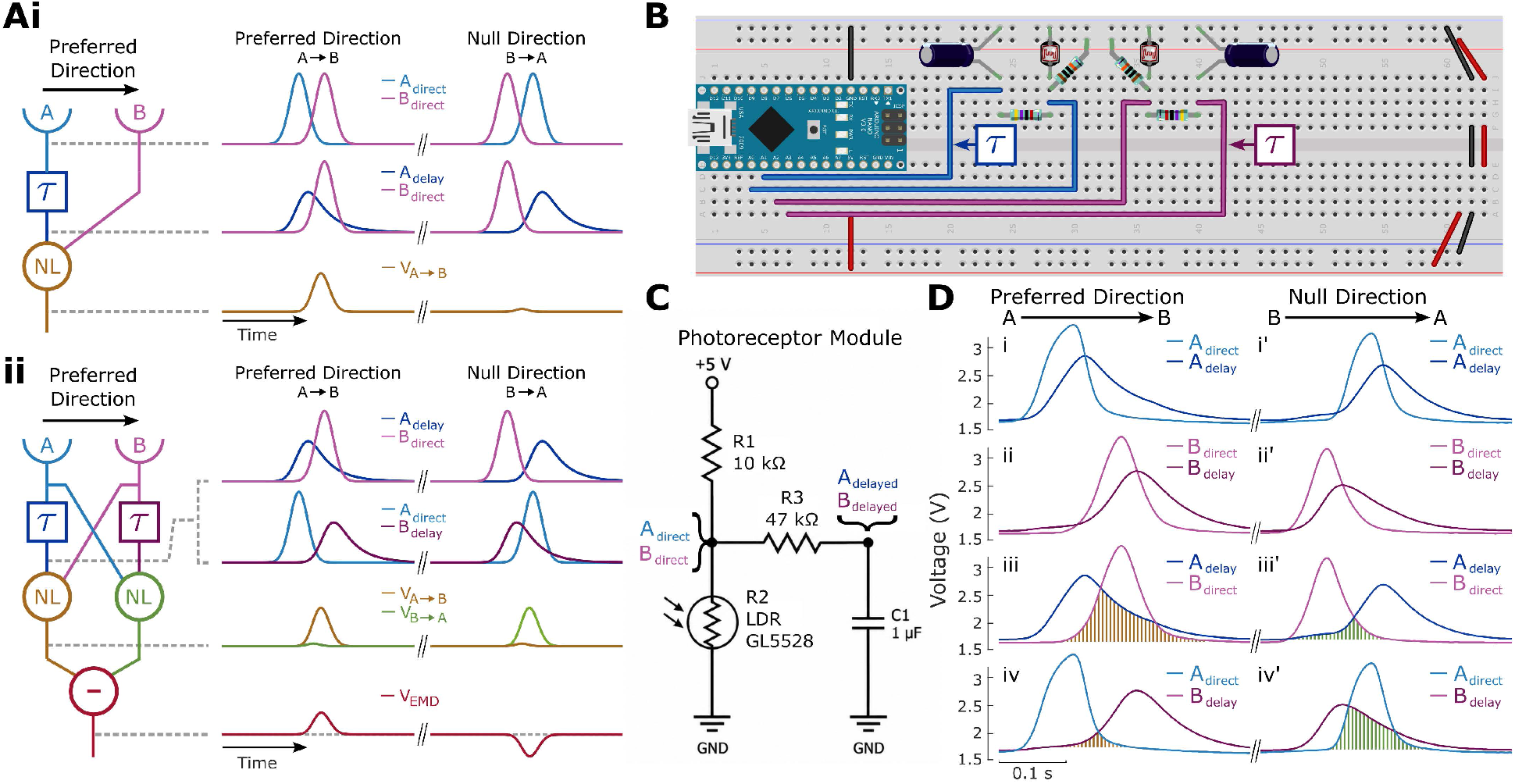
Temporal filtering and opponent elementary motion detection. **(A)** Computational architecture and responses of the elementary motion detector (EMD). **(i)** A single half-detector. Signals from two spatially separated photoreceptors, A and B, are combined after the response from A (light blue) is temporally low pass filtered with a time constant, *τ* (dark blue). During preferred-direction motion (*A* → *B*), the delayed response from A overlaps strongly with the subsequent direct response from B (magenta), producing a large output from the non-linear coincidence stage (NL), *V*_*A*→*B*_ (brown). During motion in the opposite direction (*B* → *A*), the temporal alignment between *A*_delay_ and *B*_direct_ is reduced, producing a smaller coincidence response. **(ii)** Opponent EMD containing two mirror-symmetric half-detectors. One half-detector combines *A*_delay_ with *B*_direct_ to produce *V*_*A*→*B*_ (brown), whereas the opposing branch combines *B*_delay_ with *A*_direct_ to produce *V*_*B*→*A*_ (green). Subtraction of the two branch outputs produces the opponent motion signal, *V*_EMD_ (dark red). Preferred-direction motion produces a positive response, whereas reversing stimulus direction preferentially activates the opposing branch and reverses the polarity of *V*_EMD_. **(B)** Partial breadboard implementation of the opponent EMD. Only the direct and delayed pathways are shown for each photodetector. The non-linear coincidence detection and subtraction stages are implemented in Figs. 6-7. Two GL5528 light-dependent resistors (LDRs) act as photoreceptor analogues, each providing a direct voltage signal and a temporally filtered signal through an RC low-pass filter. **(C)** Circuit diagram of the photoreceptor and temporal-filter module used in the breadboard implementation. Each GL5528 LDR forms a voltage divider with a 10 kΩ resistor to generate the direct photoreceptor voltage, *A*_direct_ or *B*_direct_. Light intensity decrements increase LDR resistance, thereby increasing *A*_direct_ or *B*_direct_. The direct signal is passed through a 47 kΩ resistor and 1 µF capacitor to form a first-order RC low-pass filter, generating the corresponding delayed signal, *A*_delay_ or *B*_delay_. **(D)** Measured direct and delayed photoreceptor responses during stimulus motion in the preferred (*A* → *B*) and null (*B* → *A*) directions. **(i, i′)** Direct and delayed responses of photoreceptor A. **(ii, ii′)** Direct and delayed responses of photoreceptor B. **(iii, iii′)** Signals entering the preferred direction (*A* → *B*) coincidence branch, *A*_delay_ and *B*_direct_. Shading indicates their temporal overlap, which is greater during preferred-direction motion. **(iv, iv′)** Signals entering the opposing *B* → *A* branch, *A*_direct_ and *B*_delay_. Their temporal overlap is greater during null-direction motion. Temporal filtering therefore converts the sequential activation of the two photoreceptors into direction-dependent differences in coincidence between the two opponent EMD half-detector branches.

A single half-detector, however, can also respond to simultaneous changes in illumination at both photoreceptors. To distinguish directional motion from such shared changes, a second, mirror-symmetric half-detector combines the filtered B signal with the instantaneous A signal (Fig. 5Aii). This opposing branch responds preferentially to motion from B to A. Subtracting its output from that of the first branch gives the opponent motion signal:

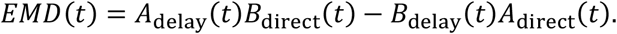

As shown in Fig. 5A, motion along the preferred direction from A to B produces a larger output from the first branch and therefore a positive opponent response (Fig. 5Aii, left traces). Reversing the direction preferentially activates the second branch, producing a negative response (Fig. 5Aii, right traces). Changes that activate both branches equally cancel during subtraction. The sign of the opponent output therefore indicates motion direction, while its magnitude reflects the imbalance between the two half-detector responses.

The breadboard implementation follows this sequence of temporal filtering, non-linear coincidence detection, and opponent subtraction (Figs. 5-7; Fig. S4). RC filters provide the temporal filtering, diode minimum-selector circuits provide a non-linear AND-like coincidence operation in place of multiplication, and an operational amplifier subtracts the two branch outputs. This circuit therefore demonstrates the computational principle of opponent motion detection.

### 2.2 Light Dependent Resistors (LDRs) as photoreceptors

In the breadboard EMD model, we can use two light-dependent resistors (LDRs; typically, cadmium sulfide photoconductive cells) positioned a short distance apart to represent neighbouring photoreceptors (Fig. 5B). The resistance of an LDR decreases with increasing illumination, allowing changes in light intensity to be converted into changes in electrical resistance. By incorporating the LDR into a voltage divider, these resistance changes can in turn be converted into changes in voltage. For two resistors connected in series, Ohm’s law gives:

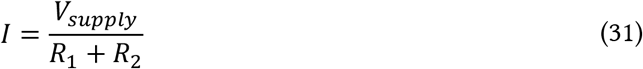

Measuring the voltage at the midpoint relative to ground is equivalent to measuring the voltage across

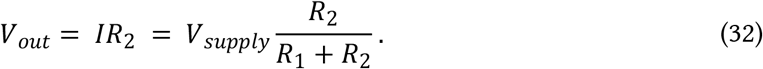

If the LDR replaces *R*_1_, increasing illumination decreases *R*_1_. Because *R*_2_ remains fixed while the total resistance *R*_1_ + *R*_2_ decreases, *V*_*out*_ increases towards *V*_*supply*_. Conversely, if the LDR replaces *R*_2_, increasing illumination decreases the fraction of the total resistance contributed by *R*_2_, causing *V*_*out*_ to decrease towards ground. Thus, the polarity of the photoreceptor voltage response can be selected simply by placing the LDR above or below the voltage-divider output.

In the present breadboard model, we will place the LDR in the *R*_2_ position of the voltage divider such that light increments decrease *V*_*out*_ (Fig. 5C). The EMD computation depends primarily on the relative timing and amplitude of the two photoreceptor signals rather than their absolute voltage polarity.

However, this arrangement provides a practical advantage because the two LDRs can be oriented towards a common light source and adjusted until their output voltages are matched to a common baseline. Matching the baseline responses of the two channels is important because any offset between them can bias the subsequent opponent comparison and produce an apparent directional preference even in the absence of motion.

### 2.3 Temporal Delay

When a moving light or dark edge passes across the two LDRs, their voltage responses occur at different times. The interval between these responses depends on the velocity of the image and the spatial separation of the detectors. This temporal ordering provides the information from which motion direction can be computed.

To compare events occurring at different times, however, the EMD must retain information about the earlier photoreceptor response until the neighbouring photoreceptor responds. In the breadboard circuit, this temporal memory is produced using an RC low-pass filter. As described in Part 1, the response time of an RC circuit is determined by its time constant, τ = RC, such that increasing either resistance or capacitance causes the output voltage to change more slowly.

Importantly, an RC filter does not produce a pure temporal shift of the input signal. Instead, it attenuates rapid voltage changes and spreads them over time. A transient photoreceptor response therefore produces a slower, more prolonged filtered response. This extended signal provides a short-term memory of the earlier visual input, allowing it to overlap with a subsequent response from the neighbouring photoreceptor.

Fig. 5B shows a partial breadboard EMD with two LDRs, each with an additional RC temporal delay pathway. Fig. 5C shows the circuit diagram for the general module used for photoreceptors A and B. When a dark object moves over the LDRs, LDR resistance increases and *V*_*out*_ increases according to Eq. 32. This *V*_*out*_ corresponds to the unfiltered photoreceptor signals A and B from Fig. 5A. To low-pass filter *V*_out_, a 47 kΩ resistor connects the photoreceptor output to a new node, while a 1 μF capacitor connects this node to ground. The filtered signals, *A*_delay_ and *B*_delay_, are measured across the corresponding capacitors (Fig. 5C). This arrangement resembles the passive membrane circuit introduced in Part 1, with the series resistor serving as the input resistance. However, the parallel membrane leak resistance is omitted to avoid attenuating the steady-state signal.

From Fig. 5D, we can see how movement in the preferred and null (or anti-preferred) direction produce different time courses of voltage changes for the A and B pathways. Along the preferred direction, the direct LDR voltage A rises before LDR voltage B (Fig. 5Di-ii). The delayed *A*_*delay*_ voltage thereby overlaps with the direct B voltage (brown shading; Fig. 5Diii). In contrast, the direct A voltage and *B*_*delay*_ overlap less (brown shading, Fig. 5Div). During null-direction motion, this pattern is reversed, with less overlap between *A*_delay_ and the direct B voltage (green shading; Fig. 5Diii′), and greater overlap between *B*_delay_ and the direct A voltage (green shading; Fig. 5Div′). This difference between preferred and null-direction responses underlies the EMD computation.

### 2.4 Non-linear Coincidence Detection

Elementary motion detection requires a non-linear operation on the direct and delayed photoreceptor signals. Simply adding the two signals would produce a large response whenever either photoreceptor was strongly activated, irrespective of whether their responses occurred at the same time. Instead, the detector must preferentially respond when the delayed signal from one photoreceptor overlaps with the direct signal from its neighbour. This temporal coincidence provides the basis for direction selectivity.

In the classical Hassenstein–Reichardt detector, coincidence is implemented by multiplication of the direct and delayed signals. Here, we use a simpler analogue approximation based on a diode minimum-selector circuit (Fig. 6A). This circuit makes use of the unidirectional conductance of a diode and can be first understood from the simplified single diode case (Fig. 6Ai). *V*_LDR_ is the voltage at the junction of the LDR and its series resistor, and therefore corresponds to the voltage across the LDR relative to ground. The output node, *V*_out_, is connected to +5 V through a 1 MΩ pull-up resistor, which tends to raise *V*_out_ towards +5 V. However, when *V*_out_ rises approximately one diode forward voltage, *V*_*F*_, above *V*_LDR_, the diode becomes forward biased and conducts. Current then flows from +5 V through the 1 MΩ resistor, through the diode, and into the LDR pathway towards ground. According to Ohm’s law, *V* = *IR*, this current produces a voltage drop across the 1 MΩ resistor, lowering *V*_out_.

**Figure 6:**
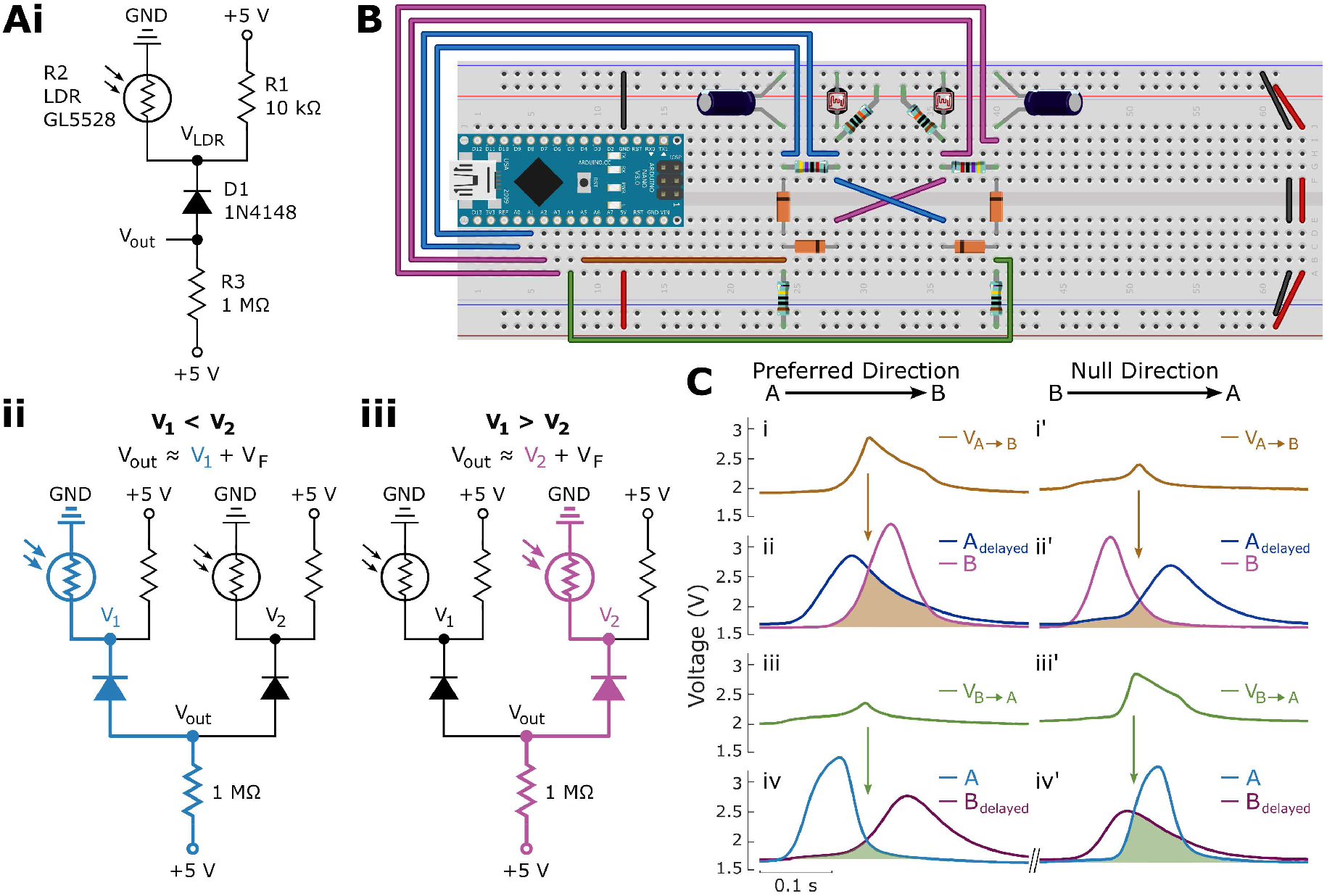
Diode-based non-linear coincidence detection in the elementary motion detector. **(A)** Principle of the diode minimum-selector circuit. **(i)** A single photoreceptor voltage, *V*_LDR_, is connected to the output node through a 1N4148 diode, while a 1 MΩ pull-up resistor connects *V*_out_ to +5 V. When *V*_out_ is approximately one diode forward voltage, *V*_*F*_, above *V*_*LDR*_, the diode becomes forward biased and conducts towards ground. This produces a voltage drop across the 1 MΩ resistor, clamping *V*_out_ to approximately *V*_LDR_ + *V*_*F*_. **(ii–iii)** With two input voltages, *V*_1_ and *V*_2_, the diode connected to the lower-voltage input experiences a greater forward bias and therefore carries more current, causing that input to dominate the output voltage. Thus, when *V*_1_ < *V*_2_, *V*_out_ ≈ *V*_1_ + *V*_*F*_, whereas when *V*_1_ > *V*_2_, *V*_out_ ≈ *V*_2_ + *V*_*F*_, where *V*_*F*_ is the diode forward voltage. The circuit therefore approximates *V*_out_ ≈ min(*V*_1_, *V*_2_) + *V*_*F*_, providing an analogue AND-like coincidence operation. Note that for illustrative simplicity, two temporally unfiltered LDRs are shown; however, in the EMD circuit, the two inputs would be one direct and one delayed signal. **(B)** Breadboard implementation of the two-channel elementary motion detector, including the paired LDR photoreceptor modules and RC delay stages from Fig. 5, together with the 1N4148 diode minimum-selector coincidence circuits. Note that there are two diode minimum-selector circuits on the breadboard, corresponding to four diodes total. In the preferred-direction (*A* → *B*) branch on the left, *V*_1_ and *V*_2_ correspond to *A*_delay_ and *B*_direct_, respectively, while in the opposing (*B* → *A*) branch on the right, they correspond to *A*_*direct*_ and *B*_*delay*_. **(C)** Example responses of the two coincidence-detector branches during motion in the preferred (*A* → *B*) and null (*B* → *A*) directions. **(i, i′)** Output of the *V*_*A*→*B*_ branch, which combines *A*_delayed_ with *B*_*direct*_. **(ii, ii′)** Corresponding direct and delayed photoreceptor signals; shaded regions indicate their temporal overlap, with the upper boundary corresponding to the minimum of the two instantaneous voltages. This minimum determines the output of the diode coincidence detector, which also includes the offset by the diode forward voltage. During preferred-direction motion (ii), *A*_delayed_ overlaps strongly with *B*_*direct*_, producing a large *V*_*A*→*B*_ response. During null-direction motion (ii’), this overlap is reduced and the response is correspondingly smaller. **(iii, iii′)** Output of the opposing *V*_*B*→*A*_ branch, combining *A*_*direct*_ with *B*_delayed_. **(iv, iv′)** Corresponding direct and delayed signals. This branch responds more strongly during *B* → *A* motion, producing the complementary direction-selective response required for opponent motion detection.

An equilibrium is reached when the diode is forward biased by approximately *V*_*F*_, such that

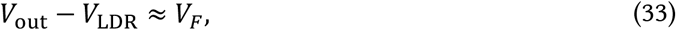

and therefore

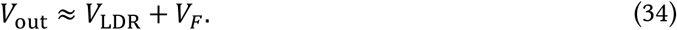

If *V*_*out*_ − *V*_*LDR*_ falls below *V*_*F*_, diode current decreases sharply, reducing the voltage drop across the 1 MΩ resistor, and allowing the pull-up resistor to raise *V*_out_ towards +5 V. Conversely, if *V*_out_ − *V*_LDR_ increases above *V*_*F*_, diode current increases sharply, producing a larger voltage drop across the 1 MΩ resistor and lowering *V*_out_. The circuit therefore settles near *V*_out_ = *V*_LDR_ + *V*_*F*_. In this way, the diode acts as a voltage clamp, limiting *V*_out_ to approximately one forward-voltage drop, *V*_*F*_, above the input signal (here, *V*_LDR_).

Adding a second diode connected to a second input, *V*_2_, introduces a second possible clamp voltage (Fig. 6Aii–iii). Each diode can constrain the output to approximately one forward-voltage drop above its respective input:

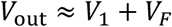

or

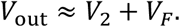

If *V*_1_ < *V*_2_, the diode connected to *V*_1_ becomes forward biased at a lower value of *V*_out_. It therefore begins conducting first and clamps the output near *V*_1_ + *V*_*F*_. At this output voltage, the diode connected to *V*_2_ is less forward biased, or becomes reverse biased, and therefore carries less current. When the two input voltages are sufficiently different, the lower-voltage input dominates the clamp; when they are similar, both diodes can contribute to conduction. Conversely, when *V*_2_ < *V*_1_, the diode connected to *V*_2_ carries more current and clamps the output near *V*_2_ + *V*_*F*_. The circuit therefore selects the lower of the two input voltages:

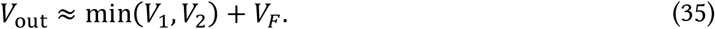

Consequently, if either input remains low, the output is held approximately one forward-voltage drop above that lower input voltage. Only when both inputs are elevated can *V*_out_ become elevated. The two-diode minimum selector therefore provides an analogue AND-like coincidence operation, producing its largest response when both input signals are simultaneously high.

In the breadboard EMD, two of these minimum-selector circuits are arranged reciprocally (Fig. 6B). One combines *A*_*delay*_ with *B*_direct_ to generate *V*_*A*→*B*_, whereas the other combines *A*_direct_ with *B*_*delay*_ to generate *V*_*B*→*A*_. During motion from A to B, photoreceptor A is activated first. Its delayed response therefore remains elevated when the direct response from B subsequently rises, producing a period of strong temporal overlap between *A*_*delay*_ and *B*_direct_ (Fig. 6Cii). Because the minimum-selector output follows the lower of these two signals, both signals must be simultaneously elevated for *V*_*A*→*B*_ to become elevated, producing the large coincidence response shown in Fig. 6Ci. For motion in the opposite direction, this temporal alignment is reduced and the *V*_*A*→*B*_ response is correspondingly smaller (Fig. 6Ci′–ii′). Conversely, motion from B to A produces stronger overlap between *B*_*delay*_ and *A*_direct_, resulting in a larger response from the opposing *V*_*B*→*A*_ branch (Fig. 6Ciii′–iv′). Thus, although the diode minimum selector is a simplified analogue approximation of the multiplicative non-linearity of the Hassenstein–Reichardt detector, it converts the temporal ordering of the two photoreceptor signals into complementary direction-selective voltage responses.

### 2.5 Opponent Subtraction and Direction Selectivity

The two complementary minimum-selector diode circuits described above generate coincidence signals for the two possible directions of motion corresponding to the nonlinear integration stages of the EMD. The minimum-selector branch on the left produces a large response when the delayed signal from photoreceptor A overlaps with the direct signal from photoreceptor B (as occurs during preferred direction motion), whereas the minimum-selector branch on the right produces a large response when the delayed signal from B overlaps with the direct signal from A (as occurs during null direction motion). A final stage is required to compare these two branches and convert their relative magnitude into a single signed motion signal.

#### 2.5.1 The Operational Amplifier

This comparison can be implemented using an operational amplifier (op-amp; Fig. 7A–B). An op-amp is a high-gain differential amplifier whose output, *V*_*out*_, depends on the difference between its non-inverting (*V*_+_) and inverting (*V*_−_) inputs:

**Figure 7:**
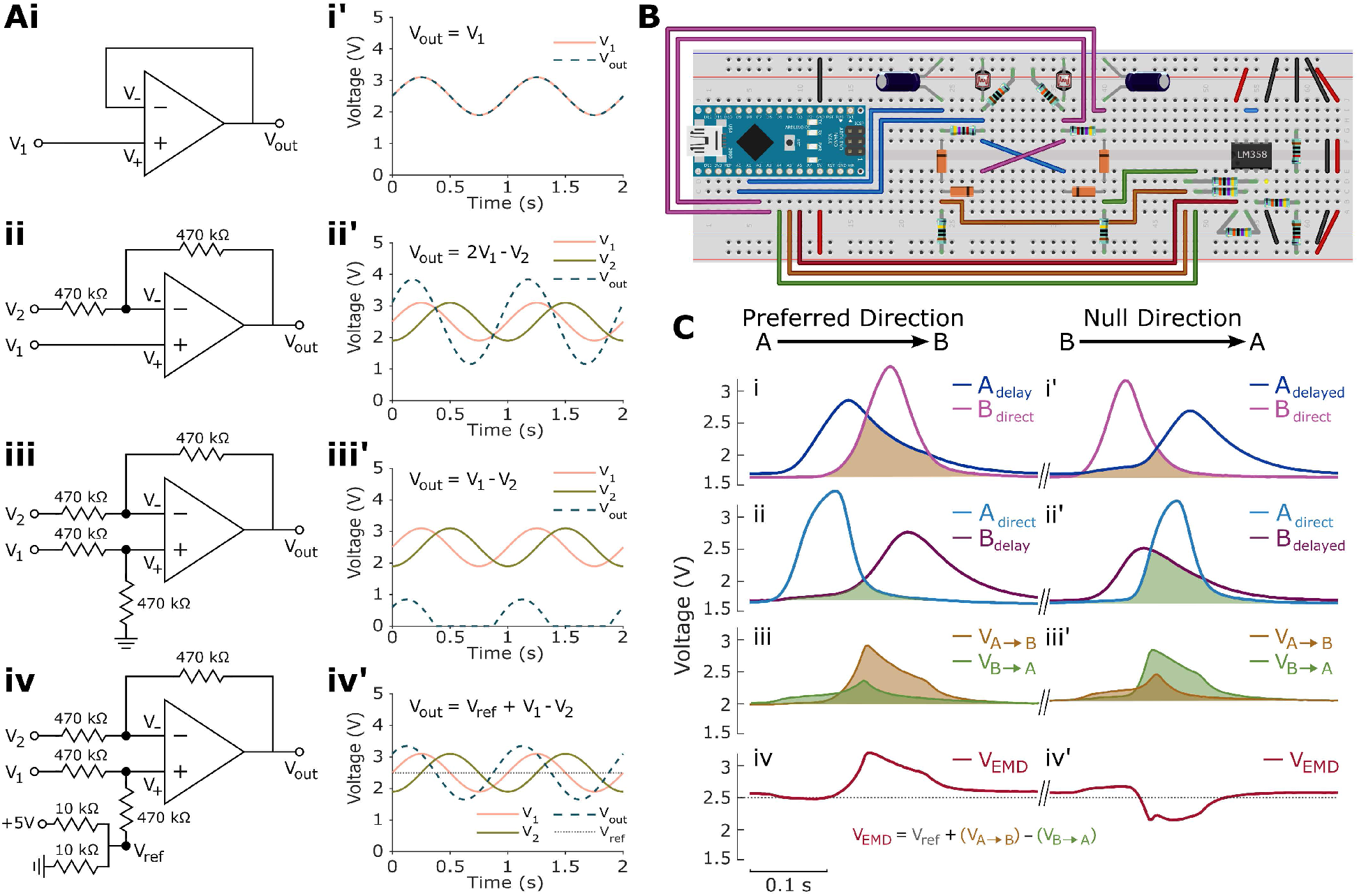
Opponent subtraction generates a signed direction-selective output from the elementary motion detector. **(A)** Stepwise construction of the op-amp subtractor. **(i)** The simplest op-amp operation is the voltage follower. The input voltage *V*_1_ is applied directly to the non-inverting input, *V*_+_, while *V*_out_ is fed back to the inverting input, *V*_−_. Negative feedback drives *V*_+_ ≈ *V*_−_, such that *V*_out_ ≈ *V*_1_. **(i′)** Simulated example sinusoidal input and corresponding follower output. **(ii)** Introduction of a second voltage, *V*_2_, at the inverting input. Equal input and feedback resistors place *V*_−_ midway between *V*_2_ and *V*_out_, while *V*_1_ remains connected directly to *V*_+_. Negative feedback therefore gives *V*_out_ ≈ 2*V*_1_ − *V*_2_. **(ii′)** Simulated example input and output voltages illustrating the unequal weighting of the two inputs. **(iii)** A matching equal-resistance voltage divider is introduced at the non-inverting input, giving *V*_+_ ≈ *V*_1_/2. Together with *V*_−_ ≈ (*V*_2_ + *V*_out_)/2, this produces the unity-gain subtraction *V*_out_ ≈ *V*_1_ − *V*_2_. **(iii′)** Simulated example subtraction of the two sinusoidal inputs. Negative values of the ideal difference cannot be represented by the single-supply 0–5 V circuit and are therefore constrained by the lower supply rail. **(iv)** To represent differences of either sign, the ground connection of the non-inverting divider is replaced by a 2.5 V reference voltage, *V*_ref_, generated from the 5 V supply. Negative feedback then gives *V*_out_ ≈ *V*_ref_ + *V*_1_ − *V*_2_. **(iv′)** The reference shifts the zero-difference level to approximately 2.5 V, allowing positive and negative differences between *V*_1_ and *V*_2_ to be represented as voltage excursions above and below the reference level. **(B)** Breadboard implementation of the complete elementary motion detector, including the LM358 op-amp used for opponent subtraction of the two dual-diode coincidence-detector outputs. Four 470 kΩ resistors form the subtractor network shown in panel Aiv, while two 10 kΩ resistors form the voltage divider used to generate the 2.5 V reference voltage. LM358 pin 1 (lower left) is the EMD output; pins 2 and 3 are the inverting and non-inverting inputs, respectively; and pins 4 and 8 are connected to ground and +5 V, respectively. The LM358 contains two independent op-amps, using pins 1–3 and 5–7. Only the first op-amp is used for opponent subtraction. The unused second op-amp is configured as a voltage follower referenced to ground to avoid leaving its inputs floating, with its non-inverting input connected to ground and its output fed back to its inverting input. **(C)** Measured signals during motion in the preferred (*A* → *B*) and null (*B* → *A*) directions. **(i, i′)** Delayed photoreceptor-A signal, *A*_delay_, and direct photoreceptor-B signal, *B*_direct_, forming the *A* → *B* coincidence pathway. Shading indicates their temporal overlap. **(ii, ii′)** Direct photoreceptor-A signal, *A*_direct_, and delayed photoreceptor-B signal, *B*_delay_, forming the opposing *B* → *A* coincidence pathway. **(iii, iii′)** Corresponding outputs of the two dual-diode minimum-selector circuits, *V*_*A*→*B*_ and *V*_*B*→*A*_. Preferred-direction motion produces the larger *V*_*A*→*B*_ response, whereas null-direction motion produces the larger *V*_*B*→*A*_ response. **(iv, iv′)** Final opponent EMD output, *V*_EMD_ ≈ *V*_ref_ + *V*_*A*→*B*_ − *V*_*B*→*A*_. The 2.5 V reference represents zero difference between the two opponent pathway outputs. Preferred-direction motion produces a positive excursion from this baseline, whereas reversing the direction of motion reverses the imbalance between the two coincidence pathways and produces an excursion of the opposite polarity. Opponent subtraction therefore converts the two complementary coincidence signals into a single signed analogue representation of motion direction.

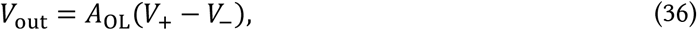

where *A*_OL_ is the open-loop gain of the op-amp. Because this gain is very large, even a small difference between the two input voltages can drive the output towards one of its supply limits. If *V*_+_ > *V*_−_, the output increases towards the positive supply rail, whereas if *V*_+_ < *V*_−_, the output decreases towards the negative or lower supply rail.

Negative feedback can be used to equalise the inputs *V*_+_ and *V*_−_, a property that allows the op-amp to be configured to perform arithmetic operations such as addition and subtraction. When *V*_*out*_ is connected back to the inverting input *V*_−_, an increase in *V*_out_ also increases *V*_−_. If *V*_+_ > *V*_−_, the op-amp increases *V*_out_ according to Eq. 36, which in turn raises *V*_−_ and reduces the difference between the two inputs. Conversely, if *V*_−_ > *V*_+_, the op-amp decreases *V*_out_ according to Eq. 36, lowering *V*_−_ and again reducing the difference. The circuit therefore reaches a stable operating point at which the two input voltages are approximately equal:

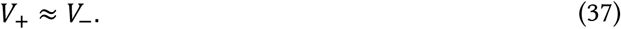

This behaviour is known as negative feedback because changes in the output act to oppose, rather than reinforce, the difference between the two input voltages.

#### 2.5.2 The Voltage Follower

The simplest application of this negative-feedback principle is the voltage follower (Fig. 7Ai). Here, an input voltage, *V*_1_, is connected directly to the non-inverting input, such that *V*_+_ = *V*_1_, while *V*_out_ is connected directly back to the inverting input, such that *V*_−_ = *V*_out_. Because negative feedback equalises *V*_+_ and *V*_−_ according to Eq. 37, the output consequently follows the input voltage:

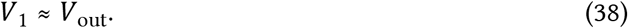

#### 2.5.3 Subtracting Input Voltages

To compare two signals, a second input voltage, *V*_2_, must also influence the inverting input (Fig. 7Aii). *V*_2_ and *V*_out_ cannot simply be connected directly to the same node because, when their voltages differ, the two voltage sources would oppose one another. Instead, each is connected to *V*_−_ through an equal resistor, *R*. Because very little current enters the op-amp input itself, current flows predominantly between *V*_2_ and *V*_out_ through the two resistors, placing *V*_−_ midway between the two voltages:

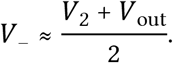

If *V*_1_ remains connected directly to the non-inverting input, then

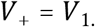

Applying the negative-feedback condition *V*_+_ ≈ *V*_−_ gives

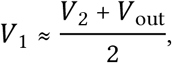

which rearranges to

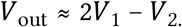

The circuit therefore already performs subtraction, but the two inputs are not weighted equally: *V*_1_ contributes with a gain of +2, whereas *V*_2_ contributes with a gain of −1. To give the two inputs equal weighting, *V*_1_ is also connected to *V*_+_ through a pair of equal resistors, with the second resistor initially connected to ground (Fig. 7Aiii). These resistors form a voltage divider, such that

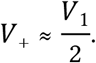

The inverting input remains

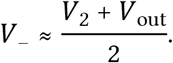

Negative feedback therefore gives

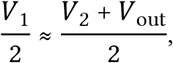

which simplifies to

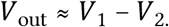

The matched resistor networks therefore produce a unity-gain subtractor: the two input voltages contribute equally but with opposite sign.

#### 2.5.4 Introducing a Reference Voltage

However, if *V*_2_ > *V*_1_, the subtraction requires a negative output voltage. With our 0–5 V circuit, it is not possible to generate negative voltages below ground. To allow differences of either sign to be represented, the ground connection in the non-inverting voltage divider is therefore replaced by a 2.5 V reference voltage, *V*_ref_, here produced from a voltage divider (Fig. 7Aiv).

In this case with equal resistors connecting *V*_1_ and *V*_*ref*_ to *V*_+_

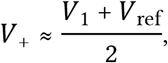

while

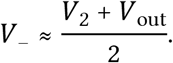

Applying *V*_+_ ≈ *V*_−_ gives

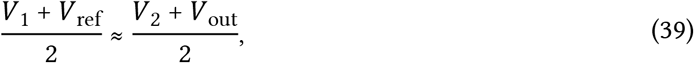

and therefore

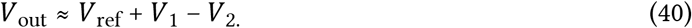

The 2.5 V reference therefore shifts the zero point of the subtraction from ground to the midpoint of the 5 V supply. Equal input voltages produce an output of approximately 2.5 V; *V*_1_ > *V*_2_ produces a positive deflection above 2.5 V, whereas *V*_2_ > *V*_1_ produces a negative deflection below 2.5 V. The circuit can therefore represent the sign of the difference between the two signals without requiring a negative power supply.

#### 2.5.5 Opponent Subtraction for the Elementary Motion Detector

In the elementary motion detector, the two input voltages to the subtractor are provided by the complementary coincidence-detector branches described in Section 2.4. We define *V*_1_ = *V*_*A*→*B*_, *V*_2_ = *V*_*B*→*A*_, and *V*_out_ = *V*_*EMD*_. Substituting these signals into the subtractor equation (Eq. 39) gives

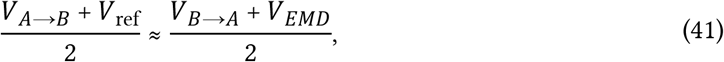

which simplifies to

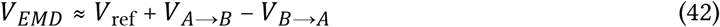

The final EMD output, *V*_*EMD*_, therefore represents the difference in activity between the two opposing motion-detector branches rather than the absolute response of either branch alone. When the two coincidence signals are equal, their difference is zero and *V*_EMD_ remains at the 2.5 V reference level. During motion from A to B, the stronger temporal coincidence between *A*_delayed_ and *B*_direct_ produces a larger *V*_*A*→*B*_ response, shifting *V*_EMD_ to one side of the 2.5 V baseline. Motion from B to A instead preferentially increases *V*_*B*→*A*_, shifting the output to the opposite side of the baseline. Thus, the magnitude of the output deflection reflects the imbalance between the two motion pathways, while its polarity indicates motion direction.

The complete subtractor is implemented on the breadboard using one of the two internal amplifiers in the LM358 dual operational amplifier (Fig. 7B–C). The two coincidence-detector outputs were connected to the non-inverting and inverting resistor networks, while a 2.5 V reference was generated by an equal-resistance voltage divider between +5 V and ground. The LM358 therefore performs the final opponent comparison entirely in analogue circuitry, converting the direct and delayed photoreceptor voltages generated earlier in the circuit into a single direction-selective output.

The measured output traces demonstrate this final operation (Fig. 7Civ). Before motion, the two opposing pathways are approximately balanced and the EMD output remains close to the 2.5 V reference. Passage of a stimulus in one direction increases one coincidence-detector branch relative to the other, producing a transient deflection of *V*_*EMD*_ away from the reference level. Reversing the direction of motion reverses which branch dominates and consequently produces a deflection of the opposite polarity. The complete breadboard circuit therefore reproduces the fundamental computational sequence of an opponent elementary motion detector: spatially separated photoreceptor signals are temporally filtered, combined through a non-linear coincidence operation, and finally subtracted to produce a signed estimate of motion direction.

## Discussion

The experiments presented here demonstrate how simple electronic components can reproduce fundamental features of neuronal dynamics and support a biologically relevant computation. The passive membrane circuit reproduces the effects of resistance, capacitance, and synaptic conductance on voltage amplitude and response kinetics (Figs. 1-2). Extending this circuit into a chain of compartments introduces spatial attenuation (Fig. 3) and temporal delay (Fig. 4), whilst combining temporal filters with non-linear coincidence detection and opponent subtraction produces direction-selective responses to visual motion (Figs. 5-7). Together, these experiments connect the electrical properties of individual components to the behaviour of an increasingly complex circuit.

The use of solderless breadboard construction complements existing equivalent-circuit exercises and electronic neuron platforms (Baden *et al*., 2018; Dabrowski *et al*., 2013; Giglia *et al*., 2019; Stradmann *et al*., 2025). Here, the emphasis is directed toward assembling and modifying the circuits using individual components, with mathematical predictions tested against measured voltages throughout. Components can be exchanged without soldering, and the same equipment can be reused across the different experiments. This flexibility allows individual exercises to be selected according to the available time and experience level of the students. For example, an introductory session could focus on predicting and measuring the effects of changing membrane capacitance, whilst a more advanced exercise could examine conductance-dependent time constants, finite-cable behaviour, or the consequences of altering the temporal filters in the motion detector.

These breadboard circuits also provide opportunities to develop practical experimental skills alongside neurophysiological understanding. In particular, constructing electronic circuits from scratch introduces students to custom electronic design, an increasingly important skill for experimental neuroscientists. Similarly, using a microcontroller to supply power and acquire voltage measurements provides an accessible introduction to microcontroller programming. By modifying circuit parameters, students also gain experience in quantitative experimentation and evaluating the agreement between theoretical predictions and physical systems.

The modular construction and flexibility of the solderless breadboard offer several directions for further exploration. Additional cable branches or inputs at different compartments could be used to investigate the influence of input location and local shunting on voltage spread. In the motion detector, varying photoreceptor separation, filter time constants, stimulus speed, and contrast would allow students to investigate how stimulus properties interact with circuit dynamics. Threshold and reset circuits could subsequently extend the passive membrane model into a leaky integrate-and-fire neuron, providing a route towards spiking networks. More broadly, breadboard neuroscience encourages students to engage creatively in designing their own experiments, testing predictions, and exploring increasingly complex principles of neuronal dynamics and computation.

## Methods and Materials

### Breadboard Components

All circuits were constructed using off-the-shelf through-hole components assembled on solderless breadboards. Arduino Nano-compatible microcontroller boards (ATmega328P) provided a nominal 5 V supply and common ground reference and were used to acquire voltage measurements through their analogue input pins, *A*_0_−*A*_7_. Table 1 details the component values and quantities used for each circuit. Resistors are all axial metal-film, 0.25 W, ±1% tolerance. Capacitors are all radial aluminium electrolytic, polarised, ±20% tolerance, unless stated otherwise.

**Table 1.** Electronic components used in the breadboard neuroscience models. Component values labelled ‘baseline’ correspond to the standard circuit configurations. Alternative resistor and capacitor values were substituted individually to investigate their effects on circuit behaviour. Quantities indicate the number required for a given component value, rather than the number installed simultaneously across all experimental conditions.

| Model | Component | Specification/Description | Quantity |
| --- | --- | --- | --- |
| Shared equipment | Arduino Nano-compatible board | 5 V, ATmega328P | 1, or 1 per model |
|  | Solderless breadboard | Standard | 1, or 1 per model |
|  | Jumper wires | Assorted lengths | As required |
|  | USB cable | Arduino power and serial connection | 1 |
| Passive membrane<br>(Fig. 1) | Membrane resistor, $R_m$ | 47 kΩ | 1 |
| | | 100 kΩ (baseline) | 1 |
| | | 220 kΩ | 1 |
| | | 470 kΩ | 1 |
| | Membrane capacitor, $C_m$ | 0.1 μF (ceramic, non-polarised) | 1 |
| | | 0.22 μF | 1 |
| | | 0.47 μF (baseline) | 1 |
| | | 1 μF | 1 |
| | Excitatory resistor, $R_{ex}$ | 330 kΩ | 1 |
|  | Momentary push button | Normally open | 1 |
| Synaptic conductances<br>(Fig. 2) | Membrane resistor, $R_m$ | 100 kΩ | 1 |
| | Membrane capacitor, $C_m$ | 0.47 μF | 1 |
| | Excitatory resistor, $R_{ex,1}$ | 47 kΩ | 1 |
| | | 100 kΩ | 1 |
| | | 220 kΩ | 1 |
| | | 330 kΩ (baseline) | 1 |
| | | 470 kΩ | 1 |
| | | 1 MΩ | 1 |
| | Additional excitatory resistor, $R_{ex,2}$ | 330 kΩ | 1 |
| | Shunting resistor, $R_{sh}$ | 47 kΩ | 1 |
| | | 100 kΩ | 1 |
| | | 220 kΩ | 1 |
| | | 330 kΩ (baseline) | 1 |
| | | 470 kΩ | 1 |
| | | 1 MΩ | 1 |
|  | Momentary push buttons | Normally open | 3 |
| Passive cable<br>(Figs. 3–4) | Membrane resistors, $R_m$ | 47 kΩ | 8 |
| | | 100 kΩ (baseline) | 8 |
| | | 220 kΩ | 8 |
| | | 470 kΩ | 8 |
| | Membrane capacitors, $C_m$ | 0.1 μF (ceramic, non-polarised) | 8 |
| | | 0.22 μF | 8 |
| | | 0.47 μF (baseline) | 8 |
| | | 1 μF | 8 |
| | Axial resistors, $R_a$ | 4.7 kΩ | 7 |
| | | 10 kΩ | 7 |
| | | 22 kΩ (baseline) | 7 |
| | | 47 kΩ | 7 |
| | Excitatory resistor, $R_{ex}$ | 22 kΩ | 1 |
|  | Momentary push button | Normally open | 1 |
| EMD<br>(Figs. 5–7) | Light-dependent resistors | GL5528 | 2 |
|  | Black heat-shrink tubing | Polyolefin, 5 mm diameter | 2 |
| | Photoreceptor voltage-divider resistors | 10 kΩ | 2 |
| | RC filter resistors | 47 kΩ | 2 |
| | RC filter capacitors | 1 μF | 2 |
|  | Signal diodes | 1N4148 | 4 |
| | Diode minimum-selector pull-up resistors | 1 MΩ | 2 |
| | Op-amp subtractor resistors | 470 kΩ | 4 |
| | Reference voltage-divider resistors | 10 kΩ | 2 |
|  | Dual operational amplifier | LM358 | 1 |

### Voltage measurements

Membrane voltages were measured using the analogue input pins of an Arduino Nano (A0 for single-compartment circuits and A0–A7 for the cable model). Voltage measurements and timestamps were transmitted over a serial connection at a baud rate of 250000 and acquired in MATLAB (MathWorks).

### Passive membrane analysis

Input onset and offset were detected automatically from the maximum and minimum of the first derivative of the voltage trace, respectively. Baseline voltage was calculated as the median voltage preceding input onset. The steady-state plateau voltage, *V*_∞_, was estimated as the median of the highest 10% of smoothed voltage samples recorded during input activation. The charging and discharging time constants (*τ*_on_ and *τ*_off_, respectively) were estimated by least-squares fits of single-exponential functions to the rising and falling phases of voltage traces. Measured *V*_∞_, *τ*_on_, and *τ*_off_ were compared with theoretical predictions derived from the circuit equations described in the main text.

### Cable model analysis

An eight-compartment passive cable was constructed from RC membrane compartments connected by axial resistances, *R*_*a*_, with excitation applied to the first compartment, *V*_0_. Input onset and offset were identified from *V*_0_ using the derivative method described above. Baseline and steady-state plateau voltages were estimated as the median voltages before input onset and towards the end of excitation, respectively, with plateau measurements restricted to at least four membrane time constants (*τ*_*m*_ = *R*_*m*_*C*_*m*_) after onset.

Measured response amplitudes were normalised to *V*_0_ and the space constant, *λ*, was estimated by least-squares fitting of the finite-cable model to the normalised voltage profiles. The theoretical space constant was derived from the steady-state difference equation for an infinite, uniform discrete cable (following Koch, 1999), yielding

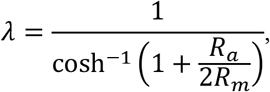

where *λ* is expressed in compartments. Electrotonic distance, *x*/*λ*, was calculated using this theoretical value. Theoretical steady-state voltage profiles were calculated by solving the system of eight linear equations describing current conservation at each compartment, incorporating membrane and axial resistances, excitatory input at *V*_0_, and a sealed distal boundary at *V*_7_.

Temporal propagation was characterised by the time to reach 50% of the response amplitude, *t*_50_, per compartment. Relative delays were calculated with respect to *V*_0_. Theoretical delays were calculated from the finite-cable model and compared with the continuous-cable prediction of *τ*_*m*_/2 per space constant for centroid delays (Agmon-Snir and Segev, 1993; Koch, 1999).

### Elementary motion detector analysis

Voltage responses from the elementary motion detector (EMD) were recorded using Arduino Nano analogue inputs *A*_0_ − *A*_6_. These comprised the direct and delayed outputs of both LDR photoreceptors (*A*_direct_, *A*_delay_, *B*_direct_, and *B*_delay_), the outputs of the two diode minimum-selector coincidence circuits (*V*_*A*→*B*_ and *V*_*B*→*A*_), and the final opponent EMD output (*V*_EMD_). Recordings were obtained during motion in both the preferred (A→B) and null (B→A) directions. Stimulus onset was identified automatically from the earliest maximum of the first derivative of the two unfiltered (direct) LDR voltage traces.

## Acknowledgements

This work was supported by a Grass Foundation Fellowship to JAS. Thanks to members of the Grass lab for helpful discussions during the development of this work.

---

**Figure S1:**
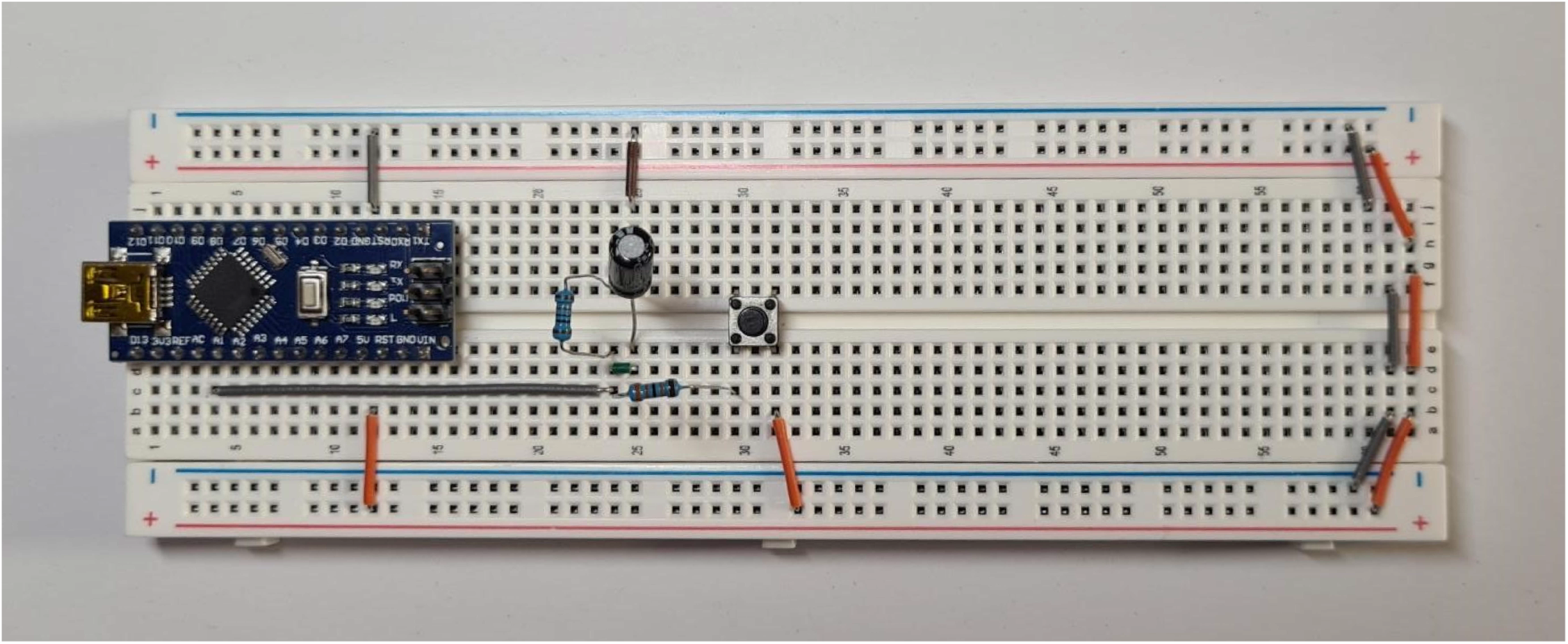
Breadboard implementation of the passive membrane circuit. Photograph showing the resistor– capacitor circuit used to model the electrical properties of a passive neuronal membrane with a single excitatory input. *R*_*m*_ = 100*kΩ, C*_*m*_ = 0.47*μF, R*_*ex*_ = 330*kΩ*.

**Figure S2:**
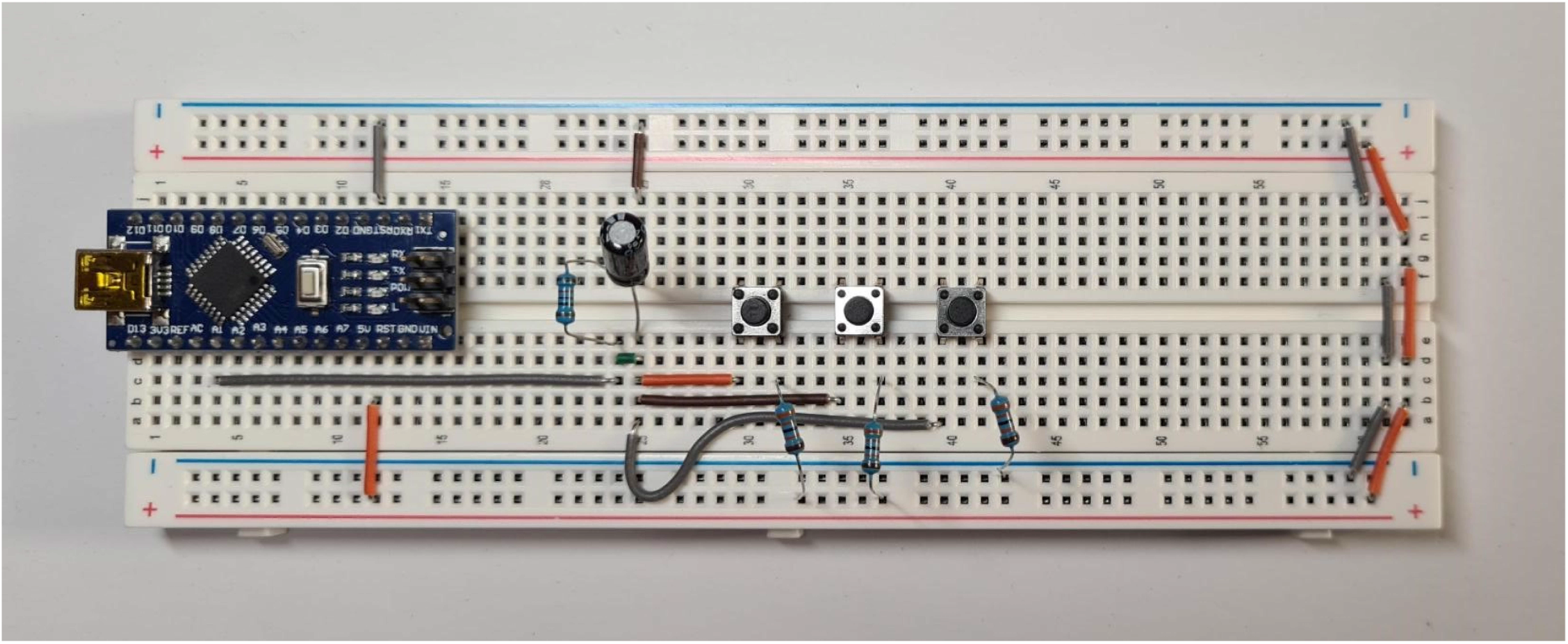
Breadboard implementation of multiple synaptic inputs. Photograph showing the passive membrane circuit extended to incorporate multiple excitatory inputs and a shunting conductance. *R*_*ex*_ = 330*kΩ, R*_*sh*_ = 330 *kΩ*.

**Figure S3:**
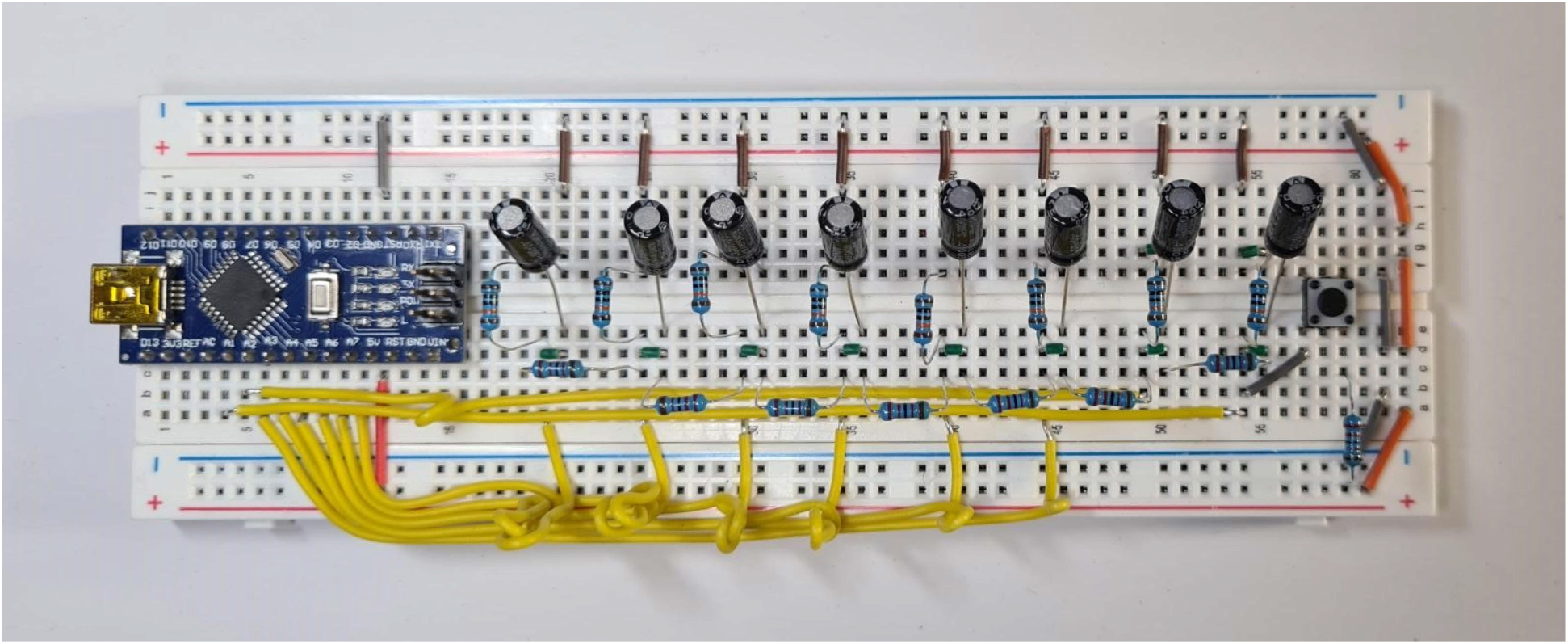
Breadboard implementation of the multi-compartment cable. Photograph showing eight passive membrane compartments connected through axial resistors to model the spatial and temporal propagation of electrical signals along a neuronal cable. *R*_*m*_ = 100 *kΩ, C*_*m*_ = 0.47 *μF, R*_*ex*_ = 22 *kΩ, R*_*a*_ = 22 *kΩ*.

**Figure S4:**
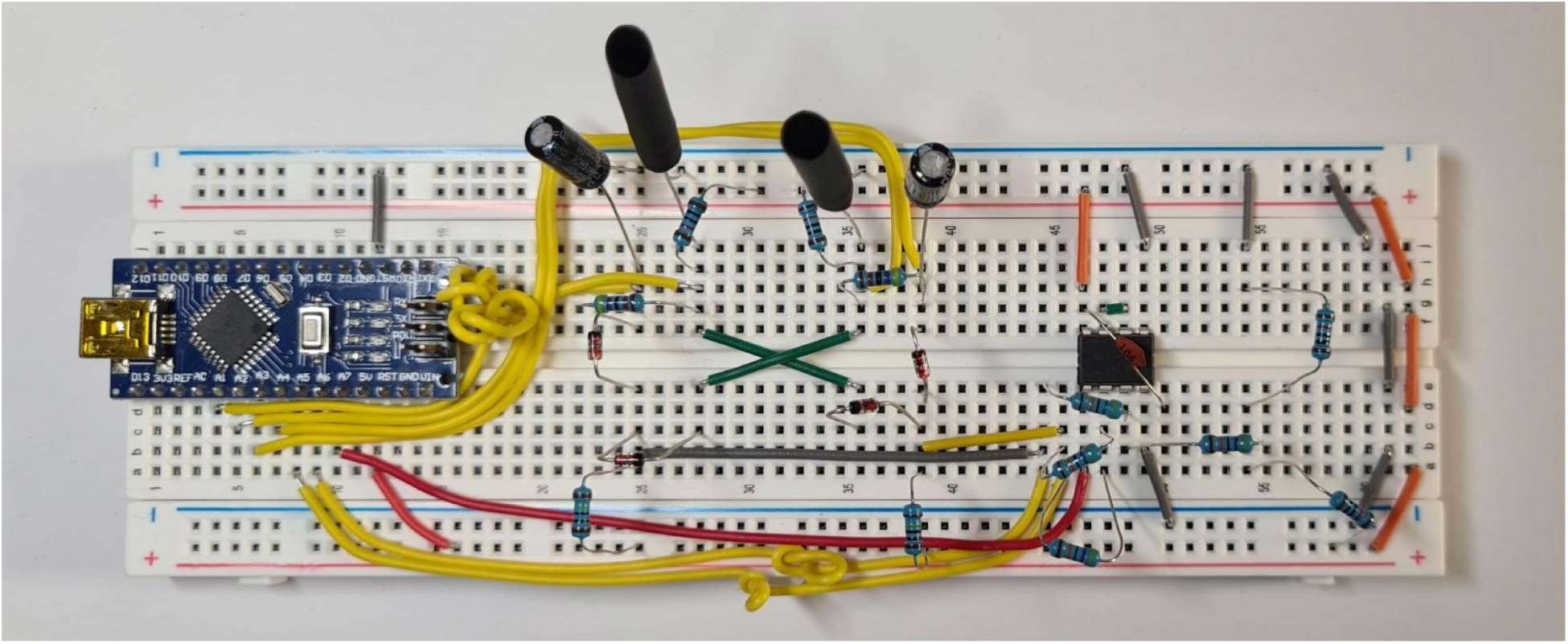
Breadboard implementation of the elementary motion detector. Photograph showing the analogue elementary motion detection circuit comprising two light-dependent resistors, RC temporal filters, diode-based coincidence circuits, and an operational-amplifier subtractor. Note that a short length of black heat-shrink tubing is mounted onto each LDR to reduce the subtended angle of light incidence, making the detectors more sensitive to the movement of small objects. Component specifications listed in Table 1.

## References

Agmon-Snir, H. and Segev, I. 1993. Signal delay and input synchronization in passive dendritic structures. J Neurophysiol, 70: 2066–2085.

Baden, T., James, B., Zimmermann, M.J.Y., Bartel, P., Grijseels, D., Euler, T., Lagnado, L. and Maravall, M. 2018. Spikeling: A low-cost hardware implementation of a spiking neuron for neuroscience teaching and outreach. PLOS Biology, 16: e2006760.

Borst, A. and Helmstaedter, M. 2015. Common circuit design in fly and mammalian motion vision. Nature Neuroscience, 18: 1067–1076.

Buchner, E. 1984. Behavioural Analysis of Spatial Vision in Insects. In: Photoreception and Vision in Invertebrates, pp. 561–621. Springer.

Curtis, H.J. and Cole, K.S. 1938. Transverse electric impedance of the squid giant axon. J Gen Physiol, 21: 757–765.

Dabrowski, K.M., Castaño, D.J. and Tartar, J.L. 2013. Basic Neuron Model Electrical Equivalent Circuit: An Undergraduate Laboratory Exercise. J Undergrad Neurosci Educ, 12: A49–A52.

Fricke, H. 1925. The electric capacity of suspensions with special reference to blood. J Gen Physiol, 9: 137–152.

Giglia, G., Crisp, K., Musotto, G., Sardo, P. and Ferraro, G. 2019. 3D Printing Neuron Equivalent Circuits: An Undergraduate Laboratory Exercise. J Undergrad Neurosci Educ, 18: T1–T8.

Hassenstein, B. and Reichardt, W. 1956. Systemtheoretische Analyse der Zeit-, Reihenfolgen-und Vorzeichenauswertung bei der Bewegungsperzeption des Rüsselkäfers Chlorophanus. Zeitschrift für Naturforschung B, 11: 513–524.

Koch, C. 1999. Biophysics of Computation: Information Processing in Single Neurons. Oxford University Press.

Marzullo, T.C. and Gage, G.J. 2012. The SpikerBox: A Low Cost, Open-Source BioAmplifier for Increasing Public Participation in Neuroscience Inquiry. PLOS ONE, 7: e30837.

Newman, M.H. and Newman, E.A. 2013. MetaNeuron: A Free Neuron Simulation Program for Teaching Cellular Neurophysiology. J Undergrad Neurosci Educ, 12: A11–A17.

Stradmann, Y., Göltz, J., Petrovici, M.A., Schemmel, J. and Billaudelle, S. 2025. Lu.i – A low-cost electronic neuron for education and outreach. Trends in Neuroscience and Education, 38: 100248.

